# A nonenzymatic dependency on inositol-requiring enzyme 1 controls cancer cell cycle progression and tumor growth

**DOI:** 10.1101/2023.11.22.567905

**Authors:** Iratxe Zuazo-Gaztelu, David Lawrence, Ioanna Oikonomidi, Scot Marsters, Ximo Pechuan-Jorge, Catarina J Gaspar, David Kan, Ehud Segal, Kevin Clark, Maureen Beresini, Marie-Gabrielle Braun, Joachim Rudolph, Zora Modrusan, Meena Choi, Wendy Sandoval, Mike Reichelt, Pekka Kujala, Suzanne van Dijk, Judith Klumperman, Avi Ashkenazi

**Author notes:** To whom correspondence should be addressed 1 DNA Way, South San Francisco, Ca, USA 94080.

## Abstract

Endoplasmic-reticulum resident inositol-requiring enzyme 1α (IRE1) supports protein homeostasis via a cytoplasmic kinase-RNase module. Known cancer dependency on IRE1 entails its enzymatic activation of the transcription factor XBP1s and of RNA decay. We discovered that some cancer cells require IRE1 but not its enzymatic activity. IRE1 knockdown, but not enzymatic inhibition or XBP1 disruption, increased DNA damage and chromosome instability while engaging the TP53 pathway and cyclin-dependent kinase inhibitors and attenuating cell cycle progression. IRE1 depletion downregulated factors involved in chromosome replication and segregation and in chromatin remodeling. Immunoelectron microscopy indicated that endogenous IRE1 can localize to the nuclear envelope. Thus, cancer cells can require IRE1 either enzymatically or nonenzymatically, with significant implications for IRE1’s biological role and therapeutic targeting.

## INTRODUCTION

Eukaryotic cells rely on a labyrinth-like membrane-surrounded organelle known as the endoplasmic reticulum (ER) to fold newly synthesized transmembrane and secreted proteins. Inositol-requiring enzyme 1α (ERN1, herein IRE1) plays a central role in maintaining ER proteostasis^1–3^. The IRE1 protein displays structural evolutionary conservation from yeast to primates, comprising an ER-lumenal domain, a single-pass transmembrane domain, and a cytoplasmic kinase-endoribonuclease module^4^. Perturbations in ER-mediated protein folding drive IRE1 enzymatic activation by direct and indirect mechanisms that lead to its homo-oligomerization, kinase-mediated trans-autophosphorylation, and allosteric RNase engagement. In turn, IRE1’s RNase triggers generation of the transcription factor XBP1s, through endomotif-based mRNA intron excision followed by nonconventional splicing^1^. Furthermore, IRE1 conducts RNase-based degradation of select mRNAs through endomotif-specific RNA cleavage—a process called regulated IRE1-dependent decay (RIDD) ^5,6^. IRE1 also performs a more promiscuous RNA decay—dubbed RIDD lacking endomotif (RIDDLE)^7^. In addition to mediating phosphate transfer between neighboring protomers within homo-oligomers^7–10^, IRE1 can trans-phosphorylate other protein substrates, such as the RNA-binding proteins Pumilio^11^ and FMRP^12^. Furthermore, IRE1 physically interacts with specific cellular proteins, including BiP^13,14^, ERDJ4^15^, PDIA6^16^, Sec61/63 translocon complexes^17^, ribosomes^18^; TRAF2^19^; non-muscle myosin^20^; and Filamin A^21,22^. In higher eukaryotes, IRE1 cooperates with two other ER resident mediators, namely PERK and ATF6^1–3^, to orchestrate the Unfolded Protein Response (UPR). The UPR helps maintain cellular proteostasis by acting to balance the ER’s protein-folding capacity with its biosynthetic load, as well as by inducing ER-associated degradation of misfolded or unfolded proteins.

Cancer cells can co-opt the UPR to mitigate ER stress caused by altered protein-folding requirements, mutations, aneuploidy, and harsh tumor microenvironments^23–26^. Cell lines derived from several cancer malignancies, including multiple myeloma (MM)^27–32^, chronic lymphocytic leukemia^33^, breast cancer ^34–37^, and colon cancer^38–41^, can exhibit a significant dependency on IRE1 for growth and survival. Studies to date have attributed this dependency to IRE1’s enzymatic activation of XBP1s^27–32,34,37,42,43^; and/or RIDD^7,44,45^. Here we show that cancer cells can possess a nonenzymatic dependency on IRE1 for growth. This finding has important potential implications for IRE1’s biological role as well as its therapeutic targeting.

## RESULTS

### The growth of some cancer cell lines requires IRE1 but not its enzymatic activity

During an investigation of human AMO1 multiple myeloma (MM) cells we were surprised to find a clear growth dependency on IRE1 without a significant requirement for XBP1. We transduced AMO1 cells with plasmids directing Doxycycline (Dox)-inducible expression of short hairpin (sh) RNAs aimed to silence either IRE1α or XBP1 mRNA transcripts. Consistent with other models^7,29^, Dox-induced expression of IRE1 shRNA (shIRE1) led to a dose-dependent depletion of both IRE1 and XBP1s, while induction of XBP1 shRNA diminished XBP1s but not IRE1 (**Fig. S1A**). As also expected, IRE1 silencing upregulated the RIDD target CD59^46^, whereas XBP1 knockdown decreased CD59 levels below baseline, consistent with a known enhancement of RIDD in the absence of XBP1^34,47,48^. Strikingly, Dox-induced IRE1 silencing strongly attenuated AMO1 cell proliferation *in vitro* and caused a marked regression of pre-established AMO1 tumor xenografts *in vivo*, whereas XBP1 silencing had little effect in both settings (**Fig. 1A and 1B**). This differential dependency on IRE1 *vs*. XBP1 suggested that AMO1 cells require IRE1 either for another enzymatic function such as RIDD—or nonenzymatically. To discern this, we blocked IRE1’s catalytic activity with an IRE1-specific small-molecule kinase inhibitor (KI)^49,50^ or RNase inhibitor (RI)^51^. In validation experiments, *in vitro* treatment of AMO1 cells with either inhibitor depleted the mRNA and protein levels of XBP1s while increasing mRNA abundance of the RIDD target DGAT2^7,52^—similar to Dox-induced IRE1 silencing, while non-targeted control shRNA (shNTC) had no effect (**Fig. S1B and S1C**). Furthermore, *in vivo* treatment of mice bearing AMO1 tumors with either IRE1 KI or RI decreased tumoral XBP1s levels and repleted DGAT2 mRNA—comparably to Dox-induced IRE1 depletion (**Fig. S1D and S1E**). Also as seen *in vitro*, XBP1 knockdown *in vivo* depleted XBP1s protein and mRNA and suppressed DGAT2 mRNA (**Fig. S1F and S1G**). Nevertheless, despite the effective blockade of IRE1’s enzymatic activity, neither IRE1 KI nor RI attenuated proliferation *in vitro* or tumor growth *in vivo* of AMO1 cells—similar to the non-targeted shRNA control, and in sharp contrast to a robust growth disruption by IRE1 silencing (**Fig. 1C and 1D**). Furthermore, the combination of either one of two different IRE1 KIs plus the IRE1 RI also did not affect AMO1 cell growth (**Fig. S1H and S1I**). Experiments with human KMS27 MM cells also revealed a dependency on IRE1 but not on XBP1 or on IRE1’s enzymatic activity (**Fig. 1E – 1H and Fig. S1J – S1M**). Moreover, *in vitro* studies with human JJN3 and L363 MM cells and with HCT116 colon carcinoma cells (**Fig. S1N – S1S**) provided additional examples of differential requirement for IRE1 *vs.* XBP1 in MM and non-MM cancer cells. In contrast, knockdown of either IRE1 or XBP1 attenuated proliferation of OPM2 MM cells (**Fig. S1T and S1U**), which were previously shown to depend on IRE1’s enzymatic activity^29^. Thus, certain cancer cell lines can possess a nonenzymatic growth dependency on IRE1.

**Figure 1.**
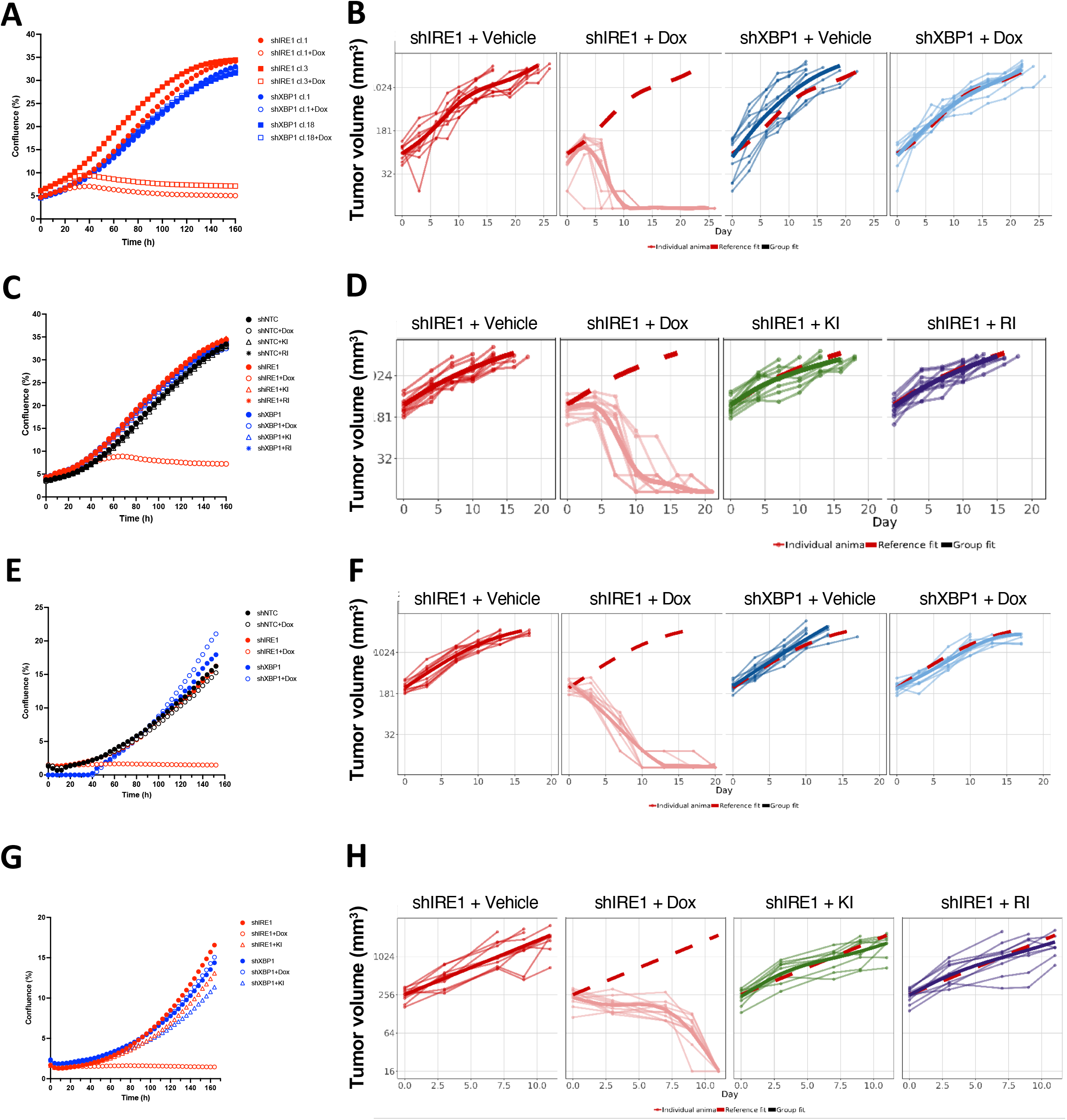
Some cancer cell lines require IRE1 but not its enzymatic activity. **(A)** Effect of IRE1 or XBP1 knockdown on *in vitro* proliferation of AMO1 cells. Cells were stably transfected with plasmids encoding doxycycline (Dox)-inducible shRNAs against either IRE1 (red) or XBP1 (blue) and grown in the absence (closed symbols) or presence (open symbols) of Dox (0.2 μg/ml). Proliferation, depicted as % confluence, was monitored by time-lapse microscopy in an Incucyte^TM^ instrument. Two independent clones (circles or squares) for each gene knockdown were characterized. **(B)** Effect of IRE1 or XBP1 knockdown on *in vivo* growth of AMO1 tumor xenografts. C.B-17 SCID mice were implanted subcutaneously with 10×10^6^ AMO1 shIRE1 cl.1 or shXBP1 cl.1 cells and allowed to form palpable tumors. Mice were then randomized into treatment groups (n=10/group) and given either vehicle (5% sucrose) or Dox (0.5 mg/ml in 5% sucrose) in drinking water *ad libitum*. Tumor growth was monitored until the last Vehicle-control animal reached endpoint (day 26). Each line represents individual mouse data, whereas the wider line represents the reference fit, corresponding to the mean growth of each grup. The wide dashed line in red represents the reference fit of shIRE1 vehicle group, embedded for comparison in all the graphs. **(C)** Effect of IRE1 or XBP1 knockdown or enzymatic IRE1 inhibition on *in vitro* proliferation of AMO1 cells. Cells stably transfected with plasmids encoding Dox-inducible shRNAs against non-targeting control (NTC, black) or IRE1 (red) or XBP1 (blue) were grown in the absence (closed symbols) or presence (open symbols) of Dox (0.2 μg/ml, circles), without or with IRE1 kinase inhibitor (KI, 3 μM, triangles) or RNase inhibitor (RI, 3 μM, asterisks). Proliferation was analyzed as in **A**. **(D)** Effect of IRE1 knockdown or enzymatic inhibition on *in vivo* growth of AMO1 tumor xenografts. C.B-17 SCID mice were implanted subcutaneously with 10×10^6^ AMO1 shIRE1 cl.1 cells and allowed to form palpable tumors. Mice were then randomized into treatment groups (n=10/group) and given either vehicle (5% sucrose) or Dox (0.5mg/ml in 5% sucrose) *ad libitum*, or treated orally bidaily with either vehicle, or IRE1 KI (250 mg/kg) or IRE1 RI (100 mg/kg). Tumor growth was monitored until the last Vehicle-control animal reached endpoint (day 19). **(E)** Effect of IRE1 or XBP1 knockdown on *in vitro* proliferation of KMS27 cells. Cells stably transfected with plasmids encoding Dox-inducible shRNAs against either IRE1 (red) or XBP1 (blue) or NTC (black) were grown, treated and analyzed as in **A**. One independent clone was characterized for each gene (cl.9 for shIRE1 and cl.13 for shXBP1). **(F)** Effect of IRE1 or XBP1 knockdown on *in vivo* growth of KMS27 tumor xenografts. C.B-17 SCID mice were implanted subcutaneously with 10×10^6^ KMS27 shIRE1 cl.9 or shXBP1 cl.13 cells and allowed to form palpable tumors. Mice were then randomized, treated and monitored as in **B** for 21 days. **(G)** Effect of IRE1 or XBP1 knockdown or enzymatic IRE1 inhibition on *in vitro* proliferation of KMS27 cells. Cells stably transfected with plasmids encoding Dox-inducible shRNAs against IRE1 (red) or XBP1 (blue) were grown in the absence (closed symbols) or presence (open symbols) of Dox (0.2 μg/ml, circles), without or with IRE1 KI (KI2, 3 μM, triangles). Proliferation was analyzed as in **A**. **(H)** Effect of IRE1 knockdown or enzymatic inhibition on *in vivo* growth of KMS27 tumor xenografts. C.B-17 SCID mice were implanted subcutaneously with 10×10^6^ KMS27 shIRE1 cl.9 cells and allowed to form palpable tumors. Mice were then randomized, treated and monitored as in **D**. Tumor growth was monitored during 12 days.

### IRE1 depletion induces cell cycle arrest independently of apoptosis

Growth deficiency due to IRE1 depletion could result from less cell proliferation or more cell death. To discern this, we stained AMO1 cells with propidium iodide (PI) to track DNA content and analyzed cell cycle progression and apoptosis by flow cytometry. Knockdown of IRE1, but not XBP1, increased the proportion of cells in the G1 phase of the cell cycle while decreasing abundance in the S and G2/M phases by 72 h (**Fig. 2A**), suggesting a failure to progress from G1 to S (G1 arrest). Kinetic studies revealed that most of the G1 accumulation occurred already by 24 h after Dox addition (**Fig. 2B**). Albeit with some delay, IRE1 knockdown also increased the incidence of cells with a subdiploid DNA content (**Fig. 2A and 2B**), suggesting some apoptosis activation. Consistent with the proliferation data (**Fig. 1**), neither XBP1 silencing nor enzymatic IRE1 inhibition disrupted cell cycle progression (**Fig. 2B and 2C**). Flow cytometry of CSFE-stained cells to track the number of completed cell divisions indicated that by 48 h a vast majority of control cells had completed two divisions, reaching generation 3 (green peak), while more IRE1-depleted cells had completed one division (orange peak) *vs.* two (green peak) (**Fig. 2D**). Confirming apoptosis induction, IRE1- but not XBP1-silenced cells showed significant cleavage of caspase-3 by 48 h, which could be blocked by the pan-caspase inhibitor QVD (**Fig. 2E**). Importantly, QVD addition during IRE1 silencing did not prevent cell cycle disruption, nor did it restore proliferation (**Fig. 2F and S2A**). Knockdown of IRE1, but not XBP1, induced G1 accumulation and apoptosis also in KMS27, JJN3, L363 and HCT116 cells (**Fig. S2B – S2E**). QVD addition blocked caspase-3 processing in KMS27 cells but did not restore proliferation and cell cycle progression by IRE1 silencing (**Fig. S2F – S2H**). Nor did QVD substantially recover proliferation of JJN3 or HCT116 cells undergoing IRE1 depletion (**Fig. S2I and S2J**). By contrast to the differential dependency of these cell lines on IRE1 vs XBP1, OPM2 cells displayed a comparable cell cycle inhibition upon silencing of either IRE1 or XBP1 (**Fig. S2K**). Flow cytometry of AMO1 cells stained with Annexin V and 7-AAD confirmed the induction of apoptosis, blocked by QVD, upon silencing of IRE1 but not inhibition of its RNase activity or knockdown of XBP1 (**Fig. S2L and S2M**). Apoptotic signaling upon ER stress can lead to caspase-mediated cleavage of IRE1^53^. To verify the uncoupling of the cell cycle inhibition from apoptosis, we generated AMO1 cells with CRISPR-mediated gene disruption of caspase-8—a protease implicated in apoptosis initiation by ER stress that is functionally suppressed by IRE1 through RIDD against DR5 mRNA^47,54–56^. As expected, caspase-8 knockout strongly dampened caspase-3/7 and apoptosis activation upon IRE1 knockdown; however, it afforded only a modest growth restoration (**Fig. S2N – S2P**). Thus, IRE1 silencing causes cell cycle inhibition independently of apoptosis activation.

**Figure 2.**
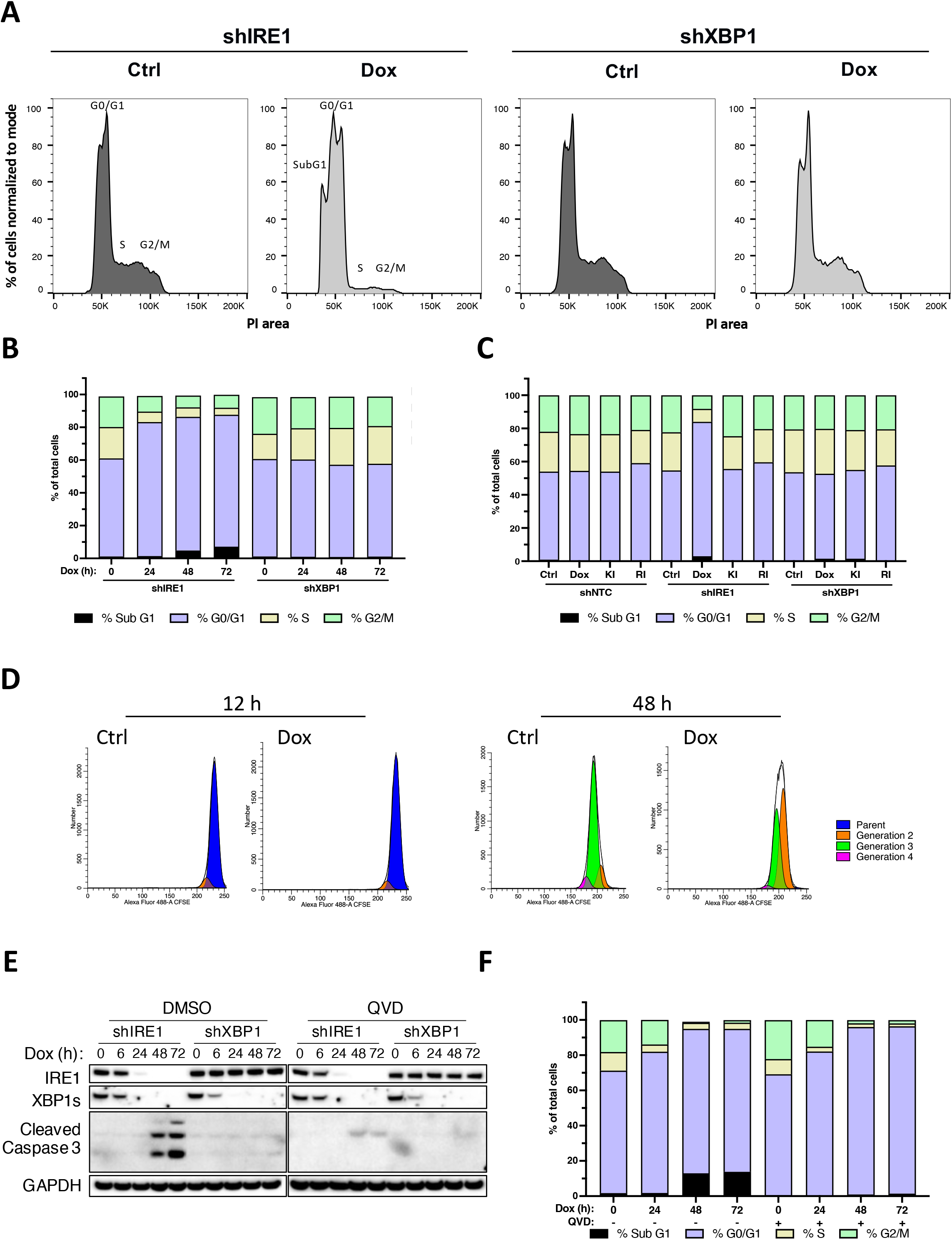
IRE1 depletion induces cell cycle arrest independently of apoptosis. **(A)** Effect of IRE1 or XBP1 knockdown on cell cycle progression. AMO1 shIRE1 cl.1 or shXBP1 cl.1 cells were incubated in the absence (dark grey) or presence (light grey) of Dox (0.2 μg/ml) for 72 h, stained with propidium iodide (PI) and analyzed by flow cytometry to determine cell cycle phases by DNA content. Sub-G1, G0/G1, S, and G2/M peaks are indicated in the shIRE1 samples. **(B)** AMO1 shIRE1 cl.1 or shXBP1 cl.1 cells were incubated with Dox (0.2 μg/ml) for the indicated time and analyzed as in **A**. **(C)** AMO1 shNTC, shIRE1 cl.1 or shXBP1 cl.1 were treated for 48 h with Dox (0.2 μg/ml), or with IRE1 kinase (KI) or RNase (RI) inhibitors at 3 μM and analyzed as in **A**. **(D)** AMO1 shIRE1 cl.1 cells were loaded with CFSE dye, incubated with Dox (0.2 μg/ml) for the indicated time, and analyzed by flow cytometry to determine generation number. **(E)** AMO1 shIRE1 cl.1 or shXBP1 cl.1 cells were treated for the indicated time with Dox (0.2 μg/ml) in the absence or presence of QVD (30 μM). Cells were lysed and analyzed by IB. **(F)** AMO1 shIRE1 cl.1 cells were treated as in **E** and analyzed by flow cytometry as in **A**.

IRE1’s enzymatic activity regulates secretory function through XBP1s as well as via RIDD^1–3^. To interrogate whether a disruption of secretion contributed to the observed dependency on IRE1, we tested whether conditioned media (CM) collected from IRE1-proficient cells could rescue the proliferation of IRE1-depleted cells. Addition of CM from IRE1-expressing AMO1 cells to naïve cells during subsequent IRE1 silencing led to a minor increase in confluence during a 7-day proliferation study; however, most of the growth inhibition persisted (**Fig. S2Q**). Conversely, media from IRE1-silenced cells did not significantly attenuate growth of IRE1-proficient cells (**Fig. S2R**). This suggested that the principal requirement for IRE1 in AMO1 cells is independent from secretion of extracellular factors.

### IRE1 silencing downregulates expression of multiple cell cycle genes

To seek further insight into the specific molecular alterations conferred by IRE1 silencing, we performed a transcriptomic analysis of AMO1 cells subjected to IRE1 *vs.* XBP1 knockdown (2 clones per gene in triplicates). We collected cells at 0, 24, 48 and 72 h after Dox addition and characterized them by bulk RNA sequencing. IRE1 or XBP1 silencing showed expected effects on the IRE1 and XBP1s proteins and their encoding mRNAs, as well as on the XBP1s transcriptional targets SEC61A and SYVN1 and the RIDD mRNA targets DGAT2 and BCAM (**Fig. S3A – S3C**). Gene-set enrichment analysis (GSEA) indicated two major ontological categories that were most notably downregulated by IRE1 silencing: Protein Processing in the ER, and Cell Cycle Control (**Fig. 3A**). Further dissection of the Cell Cycle category revealed markedly attenuated expression of genes involved in mediating the S as well as G2/M phases (**Fig. 3B – 3C** and **Fig. S3D – S3E**). In keeping with the flow cytometry data (**Fig. 2B**), many of the transcriptional changes related to cell cycle control occurred already within 24 h of Dox addition. Indeed, Hallmark GSEA further revealed significant <u>down</u>regulation of E2F, Myc, G2/M, and mitotic spindle targets (**Fig. 3D**). Gene ontology (GO) analysis further indicated depletion of mRNAs linked to DNA Replication, Chromosomal Region, Chromosome Centromeric Region, and Chromosome Segregation (**Fig. 3E**). Thus, silencing of IRE1, but not of XBP1, decreased the transcription of many cell cycle genes involved in chromosome replication, organization and segregation.

**Figure 3.**
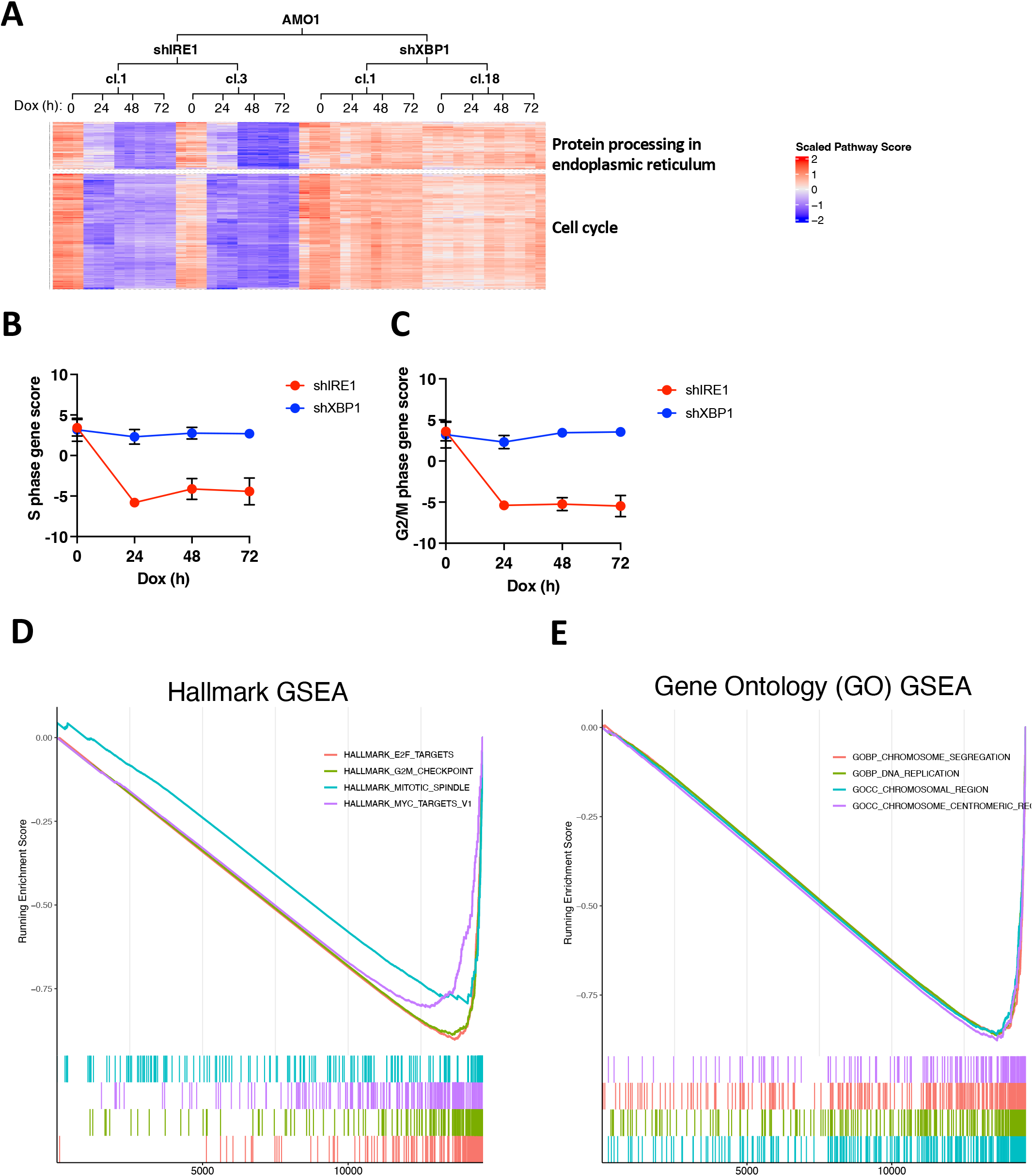
IRE1 silencing downregulates expression of multiple cell cycle genes. **(A)** Effect of IRE1 or XBP1 knockdown on mRNA expression of genes involved in Protein Processing in the ER or in Cell Cycle control. AMO1 shIRE1 cl.1 or cl.3 cells and shXBP1 cl.1 and cl.18 cells were incubated for the indicated time with Dox (0.2 μg /ml) in triplicates and subjected to bulk RNA sequencing. Shown are heat maps indicating expression of distinctly regulated mRNAs encoded by KEGG pathways gene sets involved in the indicated cellular functions by Gene Set Enrichment Analysis (GSEA) **(B)** Effect of IRE1 or XBP1 knockdown on mRNA expression of genes involved in the S phase of the cell cycle, calculated as S phase gene score. Depiction of S phase gene score for samples in **A**. **(C)** Effect of IRE1 or XBP1 knockdown on mRNA expression of genes involved in the G2 and M phases of the cell cycle, calculated as G2/M phase gene score. Depiction of G2/M phase gene score for samples in **A**. **(D)** RNAseq data from mRNAs specifically regulated upon IRE1 knockdown after 24 h, but not upon XBP1s, were queried by GSEA. Enrichment plots from the most significantly downregulated Hallmark. **(E)** Enrichment plots as in **D** for Gene Ontology (GO) gene sets.

### IRE1 depletion upregulates the p53 pathway and specific CDK inhibitors

Amongst gene-sets that were significantly <u>up</u>regulated by IRE1 silencing was the TP53 (p53) tumor suppressor pathway, which showed elevated expression of most of its target-gene clusters (**Fig. 4A** and **S4A**). To obtain additional insights, we performed a comparative proteomic analysis of AMO1 cells during knockdown of IRE1 *vs.* XBP1 (2 clones each). We collected cells at 0 and 24 h after Dox addition and characterized proteolytic digests of cell lysates by liquid chromatography-mass spectrometry (LC-MS/MS). Several proteins involved in cell cycle control displayed selective downregulation upon IRE1 silencing, including KIF22, UHRF1, KITH and KIFC1 (p < 0.05). Additional proteins that did not reach statistical significance but were notable nonetheless included AURKB, SAS6, TPX2, CCNA2, CCNB1, NUF2 and CDCA5 (**Fig. 4B**). Conversely, the cyclin dependent kinase (CDK) inhibitor CDKN1A (p21) showed significant mRNA and protein upregulation (**Fig. 4B and 4C**). Immunoblot (IB) analysis indicated upregulation of the p53 protein as early as 6 h after Dox addition, while p21 showed accumulation by 12 h (**Fig. 4D**). The gain in p53 was accompanied by its phosphorylation on Ser15 (**Fig. 4E**), indicating functional engagement^57^. RT-qPCR analysis further revealed that the upregulation of p53 was non-transcriptional, whereas p21 induction was evident both at the mRNA and protein levels (**Fig. 4D and 4F**), consistent with the known transcriptional activation of p21 by p53^58^. The proteomic data also suggested that IRE1 silencing upregulated the CDK inhibitor CDKN1B/p27 (**Fig. S4B**). This upregulation appeared to be non-transcriptional based on RNAseq and RT-qPCR analysis (**Fig. S4C and S4D)**, and was confirmed by IB at the protein level for both AMO1 and KMS27 cells (**Fig. S4E**). Because DNA damage often drives p53 activation, we interrogated specific markers of the DNA damage response (DDR) that act upstream to p53 in response to DNA double-strand breaks^59^. Indeed, knockdown of IRE1 but not XBP1 induced time-dependent upregulation and phosphorylation of histone H2AX (**Fig. S4F**); it also increased 53BP1 levels in concert with engagement of p53 (**Fig. S4G**), suggesting that IRE1 depletion augments DNA damage.

**Figure 4.**
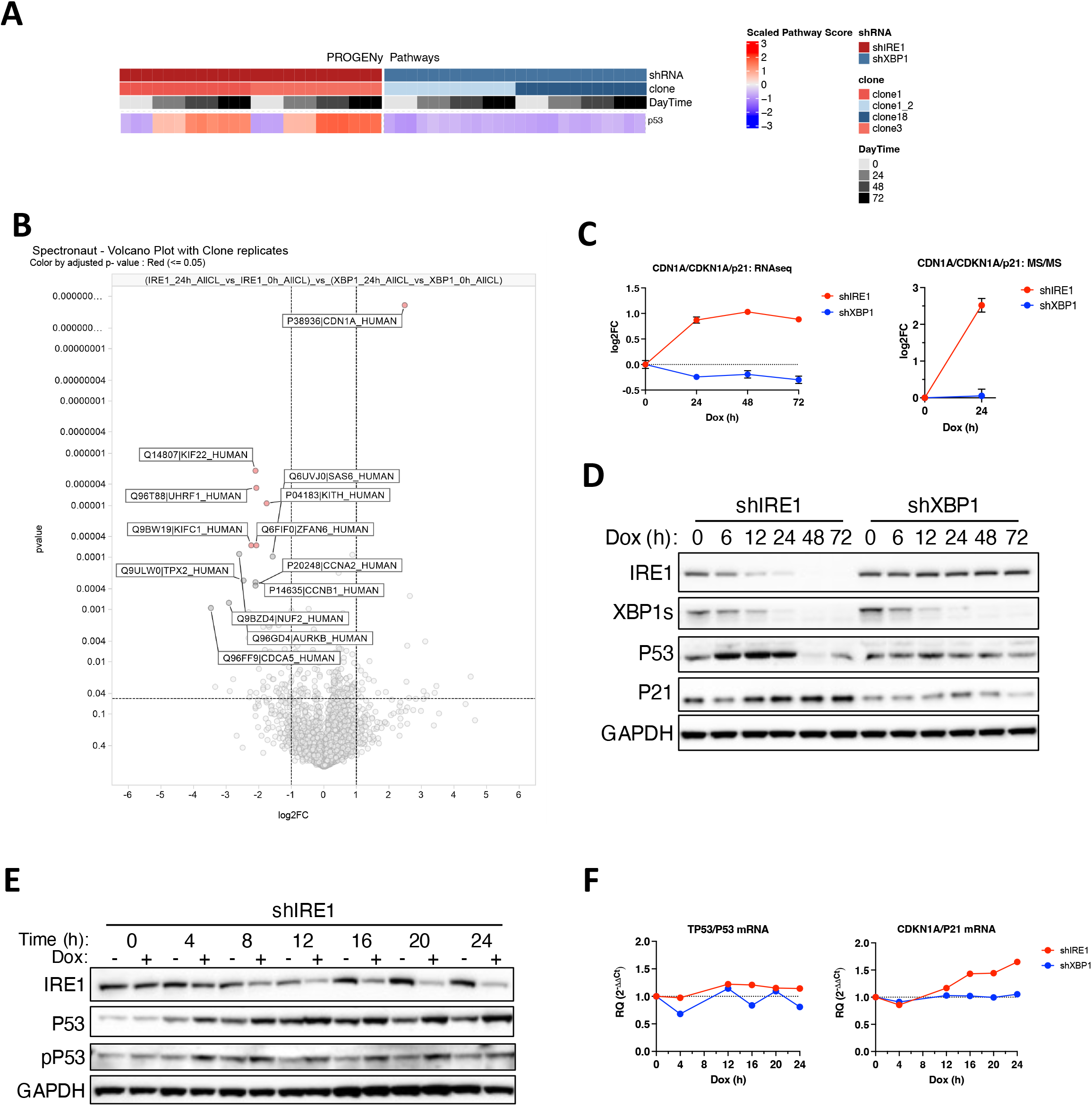
IRE1 depletion upregulates the p53 pathway and specific CDK inhibitors. **(A)** Progeny pathway analysis of the effect of IRE1 or XBP1 knockdown on mRNA expression. Heatmap depicting the top signaling pathways that are upregulated by IRE1 knockdown and downregulated upon XBP1s knockdown for samples depicted in Fig. 3A. The p53 signaling pathway (red square) stands out to be already upregulated upon IRE1 knockdown by 24 hours. **(B)** Effect of IRE1 or XBP1 knockdown on the proteome of AMO1 cells. AMO1 shIRE1 cl.1 or cl.3 cells and shXBP1 cl.1 and cl.18 cells were incubated for 24 h with Dox (0.2 μg /ml) in triplicates and analyzed for protein abundance by MS/MS. Depicted is a volcano plot showing log2 fold change (FC) in protein abundance as a function of p value, comparing t=24 h over t=0 for shIRE1 vs shXBP1. The most significantly altered proteins are labeled. Proteins marked in red show an adjusted p.value < 0.05. **(C)** Effect of IRE1 or XBP1 knockdown on CDKN1A/p21 mRNA (left) and protein (right) levels. Data depicted are from RNAseq and proteomics analyses described in Fig. 3A and 4B, respectively. **(D)** Effect of IRE1 or XBP1 knockdown on p53 and p21 protein levels. AMO1 shIRE1 cl.1 or shXBP1 cl.1 cells were incubated for the indicated time with Dox (0.2 μg /ml) and analyzed by IB. **(E)** Effect of IRE1 knockdown on p53 and phospho-p53 (pP53) levels. AMO1 shIRE1 cl.1 were incubated for the indicated time in the absence or presence of Dox (0.2 μg /ml) and analyzed by IB. **(F)** Effect of IRE1 or XBP1 knockdown on p53 and p21 mRNA levels. AMO1 shIRE1 cl.1 or shXBP1 cl.1 cells were incubated for the indicated time with Dox (0.2 μg /ml) and analyzed by RT-qPCR.

### IRE1 silencing increases chromosome instability

To investigate the effect of IRE1 silencing on chromosomal integrity, we examined Hoechst-stained AMO1 cells for the presence of micronuclei by fluorescence microscopy. Over 24 – 48 h of Dox addition, IRE1 silencing increased the relative frequency of micronuclei as compared to XBP1 knockdown (**Fig. 5A and 5B**), suggesting exacerbated chromosomal instability in IRE1-depleted cells^60^. In concert, IRE1 silencing also decreased the relative frequency of mitotic cells at 24 and 48 h and increased the incidence of apoptotic nuclei at 48 h (**Fig. 5A and 5B**). Treatment with QVD did not significantly alter the frequency of micronuclei or mitotic figures (**Fig. 5C**), confirming the increase in micronuclei caused by IRE1 knockdown and suggesting an independence of this feature from caspase activation. Furthermore, IRE1 depletion in the context of QVD treatment led to a significant decrease in DNA content (**Fig. 5D**), suggesting a specific loss during cell division. Together, these results indicate that IRE1 silencing promotes DNA damage and chromosome instability, at least in this context of aneuploidy. In keeping, GSEA of the proteomics data indicated that ontologies most significantly impacted by IRE1 silencing included: Metaphase-Plate Congregation, Mitotic Sister Chromatid Segregation, and Mitotic Cell Cycle Arrest (**Fig. S5A**). Additional notable ontologies included Cyclin-Dependent Protein Ser/Thr Kinase Inhibitor Activity and Regulation of Cell Cycle G1/S Transition. Moreover, a majority of the proteins modulated by IRE1 silencing were annotated as nuclear, whereas proteins affected by XBP1 knockdown were mainly non-nuclear (**Fig. S5B**).

**Figure 5.**
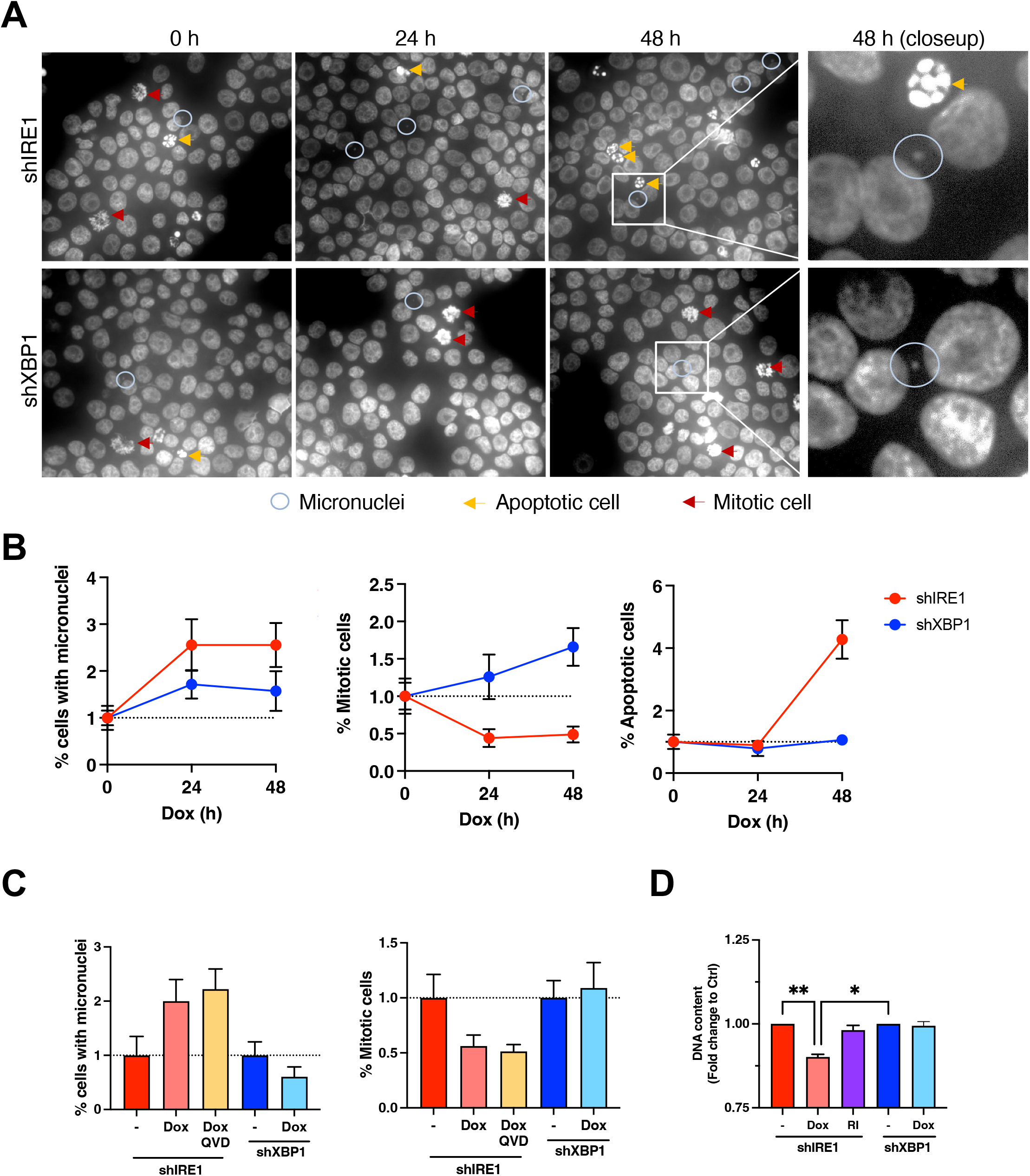
IRE1 depletion increases chromosome instability. **(A)** Effect of IRE1 or XBP1 knockdown on frequency of micronuclei. AMO1 shIRE1 cl.1 and shXBP1 cl.1 were treated for the indicated time with Dox (0.2 μg /ml), stained with Hoechst DNA dye and analyzed by fluorescence microscopy. Representative images of at least 10 fields per condition are shown. Blue circles indicate examples of cells with micronuclei (blue). Arrows indicate mitotic cells (red) or apoptotic cells (white). **(B)** Quantification of the results exemplified in **A** per total amount of cells per field. 10 fields per condition with at least 100 cells were counted. Fold change to untreated controls for each cell line is depicted. **(C)** Effect of QVD on micronuclei and mitotic frequencies. AMO1 shIRE1 cl.1 and shXBP1 cl.1 cells were incubated for 24 h with Dox (0.2 μg /ml) in the absence or presence of QVD (30 μM) and analyzed as in **B**. **(D)** Effect of IRE1 or XBP1 knockdown on DNA content. AMO1 shIRE1 cl.1 and shXBP1 cl.1 were treated for 24 h with QVD (30 μM) in the presence or absence of Dox (0.2 μg /ml) or RI (3 uM), stained with the DNA dye Hoechst, and analyzed by flow cytometry. Fold change to untreated controls of the DNA content per total amount of live cells is depicted. Mann-Whitney non-parametric U test.

### IRE1 depletion decreases UHRF1, DNA and H3 methylation, and heterochromatin

One of the most significantly downregulated proteins in IRE1-depleted cells was the DNA methylation reader UHRF1 (**Fig. 4B**)—a multidomain protein that promotes DNA methylation and chromatin modification, coordinating cell cycle control in several cancers^61–63^. Both the transcriptomic and proteomic data indicated a decrease of approximately 4 – 5-fold in UHRF1 levels by 24 h of IRE1 silencing (**Fig. 6A**). IB analysis confirmed a substantial downregulation of UHRF1 protein in response to IRE1 silencing in both AMO1 and KMS27 cells (**Fig. 6B**). To further examine UHRF1 modulation, we interrogated tumor samples from AMO1 xenografts. Dox-induced IRE1 silencing but not enzymatic IRE1 inhibition led to a marked downregulation of UHRF1, in concert with upregulation of p53, p21 and p27 (**Fig. S6A**). Similarly, tumor samples from KMS27 xenografts showed strong UHRF1 downregulation and p27 upregulation upon IRE1 knockdown, but not under enzymatic IRE1 inhibition (**Fig. S6B**).

**Figure 6.**
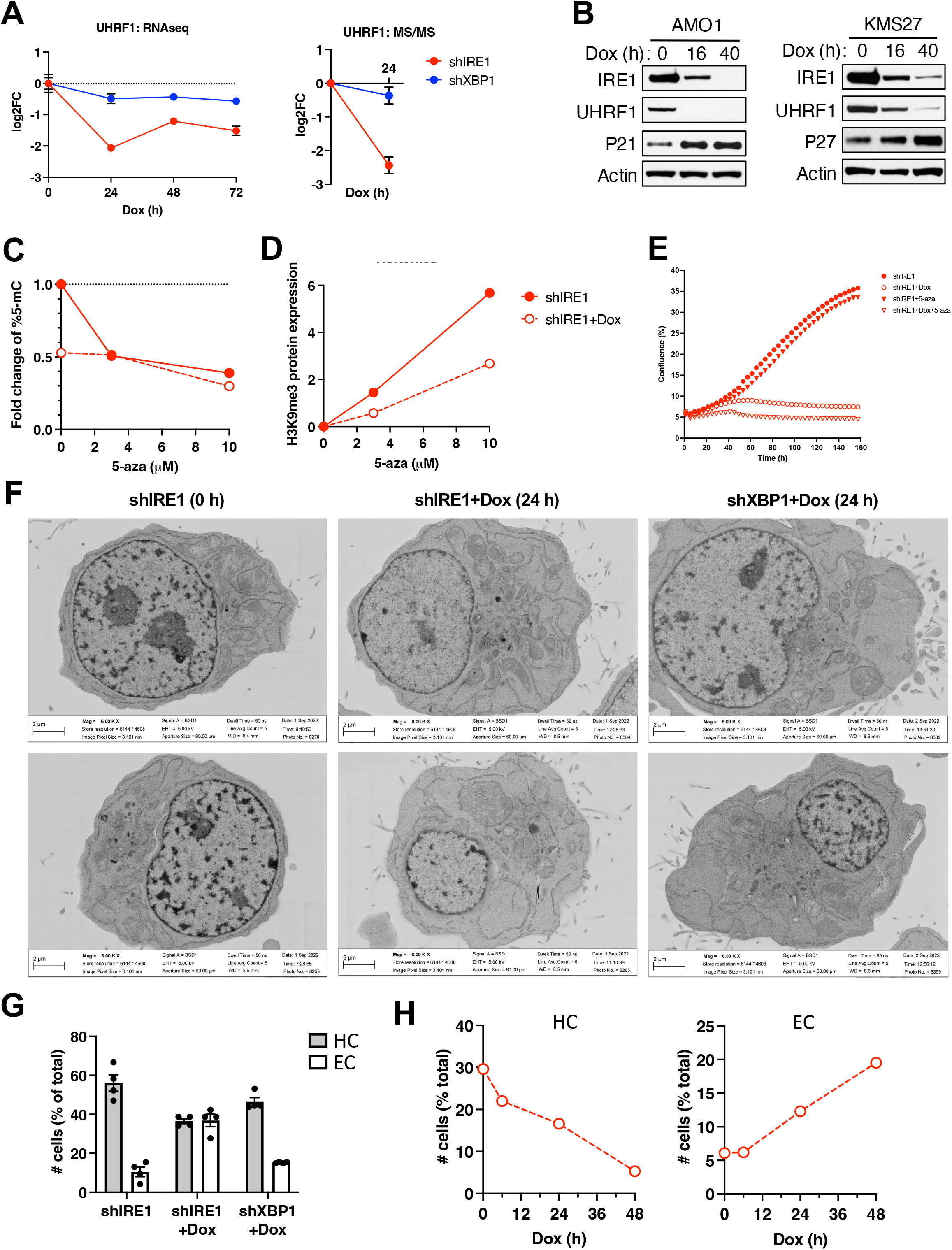
IRE1 silencing decreases UHRF1, DNA and H3 methylation. **(A)** Effect of IRE1 or XBP1 knockdown on UHRF1 mRNA (left) and protein (right) levels. Data depicted are from RNAseq and proteomics analyses described in Fig. 3A and 4B, respectively. **(B)** Effect of IRE1 knockdown on UHRF1 protein levels. AMO1 shIRE1 cl.1 and KMS27 shIRE1 cl.9 cells were incubated for the indicated time with Dox (0.2 μg /ml) and analyzed by IB. Data depicted are from the same experiment as in **Fig. S4E**. **(C)** Effect of IRE1 knockdown on DNA methylation. AMO1 shIRE1 cl.1 cells were incubated for 24 h in the absence (open symbols) or presence (closed symbols) of Dox (0.2 μg /ml) with the indicated concentration of 5-azacytidine (5-aza). Global DNA methylation levels were quantified by 5-methylcytosine (5-mc) ELISA, normalized by gDNA amount and shown as fold change to the untreated control. **(D)** Effect of IRE1 knockdown on H3K9 trimethylation. AMO1 shIRE1 cl.1 cells were incubated for 24 h in the absence (open symbols) or presence (closed symbols) of Dox (0.2 μg /ml) with the indicated concentration of 5-aza. H3K9me3 levels were quantified by IB against H3K9me3 as compared to total H3 levels (IB images are shown in **Fig. S6C**; the 5-aza concentrations of 0, 3 and 10 μM are depicted in the graph). **(E)** Effect of IRE1 knockdown and 5-aza treatment on proliferation. AMO1 shIRE1 cl.1 cells were incubated for the indicated time in the absence (open symbols) or presence (closed symbols) of Dox (0.2 μg /ml), without (circles) or with (triangles) 5-aza (10 μM). Proliferation, depicted as % confluence, was monitored by time-lapse microscopy in an Incucyte^TM^ instrument **(F)** Effect of IRE1 or XBP1 knockdown on heterochromatin density. AMO1 shIRE1 cl.1 and shXBP1 cl.1 cells were incubated for the indicated time with Dox (0.2 μg /ml). Cells were then analyzed by electron microscopy (EM). Images depict 6000X magnification. **(G)** AMO1 cells as in F were analyzed by EM to determine the frequency of nuclei with electron-dense heterochromatin (HC) or dispersed euchromatin (EC). For each sample 4 fields were analyzed at 1000X magnification with approximately 30 cells per field. **(H)** An independent unbiased analysis was performed at a different institution (Utrecht) to determine HC and EC frequencies in AMO1 shIRE1 cl.1 cells incubated for the indicated time with Dox (0.2 μg /ml). For each sample 164-196 cells were examined.

UHRF1 plays a key role in maintenance of DNA methylation and in chromatin modification during DNA replication^61–63^. We therefore examined the effect of IRE1 silencing on DNA methylation. As a control, we used the cytosine analog 5-azacytidine (5-aza), which displaces cytosine within newly synthesized DNA thereby blocking methylation. A 5-methylcytosine ELISA indicated that 5-aza treatment maximally attenuated global DNA methylation by approximately 60%, while IRE1 knockdown alone decreased this modification by about 50% (**Fig. 6C**). In presence of higher 5-aza concentrations, IRE1 deletion did not further decrease DNA methylation, suggesting convergence of the two interventions. We also interrogated histone H3 methylation by IB against H3 and H3K9me3. Although H3K9 was not detected at baseline, treatment with 5-aza increased relative H3K9me3 levels in a dose-dependent manner (**Fig. 6D and S6C**), consistent with earlier findings^64^; IRE1 knockdown substantially attenuated this elevation in H3K9me3. Furthermore, whereas 5-aza alone had a weak anti-proliferative effect on AMO1 cells, its combination with silencing of IRE1 but not XBP1 led to a more complete growth inhibition than did IRE1 disruption by itself (**Fig. 6E and S6D**). Since 5-aza inhibits DNA methylation while increasing H3K9me3 marks, the disruption of both events by IRE1 depletion may enable stronger growth inhibition.

To further interrogate the impact of IRE1 knockdown on chromatin, we examined nuclear morphological features of AMO1 cells by electron microscopy (EM). Based on electron density and distribution features, we could readily discern heterochromatin, evident from clusters of electron-dense material; nucleoli; and euchromatin, indicated by dispersed patches of relatively electron-lucent material. At 24 h after Dox addition, IRE1-silenced cells showed a marked reduction in heterochromatin, whereas XBP1-depleted cells showed similar heterochromatin density to that of baseline controls (**Fig. 6F and S6E**). Quantification of multiple fields (excluding dead cells and cells with no visible nuclei) indicated that IRE1 knockdown equalized the proportion of cells displaying dense heterochromatin or dispersed euchromatin; in contrast, baseline control and XBP1 silencing resulted in higher ratios of cells with heterochromatin vs. euchromatin (**Fig. 6G**). We performed an independent EM study at a different laboratory (University of Utrecht), in which the investigators were unaware of the latter results. Their study yielded similar data, indicating a decrease in cells with heterochromatin and a corresponding increase in cells with euchromatin in response to IRE1 silencing (**Fig. 6H and Fig. S6F**). It further provided evidence of decreased mitotic figures and increased apoptotic figures within 24 – 48 h of Dox addition (**Fig. S6G – S6I**). Thus, IRE1 depletion attenuates UHRF1 expression, DNA and H3 methylation, and heterochromatin abundance.

### IRE1 silencing downregulates multiple cell cycle proteins

IB analysis of multiple cell cycle proteins confirmed that their alteration by IRE1 knockdown was non-enzymatic (**Fig. S7A** and **S7B**). To further decipher these molecular changes, we synchronized AMO1 cells in the G1 phase by treatment with an established CDK4/6 inhibitor^65^. We then washed out the inhibitor to allow the cells to resume cycling more synchronously. At t=0 after CDK4/6 inhibitor washout, over 80% of control cells appeared in G1 (**Fig. 7A and S7C**)—as compared to the 60% we detected without synchronization (**Fig. 2B**). Cells advanced into S phase by 8 h, entering G2 and M by 12 – 20 h (**Fig. 7A and S7C**). IRE1 knockdown during the synchronization period increased the proportion of cells in G1 to approximately 90% at t=0, and markedly impaired progression from G1 into S and further into G2 and M. Relative to controls, IRE1-silenced cells showed an upregulation of γH2AX and p21 at t=0, possibly due to the depletion of IRE1 during synchronization, and an attenuated accumulation of UHRF1 over time. IRE1 knockdown also altered the kinetic profiles of several proteins involved in G1/S, including CCNA2, Rb, pRb, E2F2 and TK1; as well as proteins involved in G2/M, namely CCNB1, CDC20, CDCA5, CDCA8, AURKB, Geminin, Securin, KIF22 and pH3.

**Figure 7.**
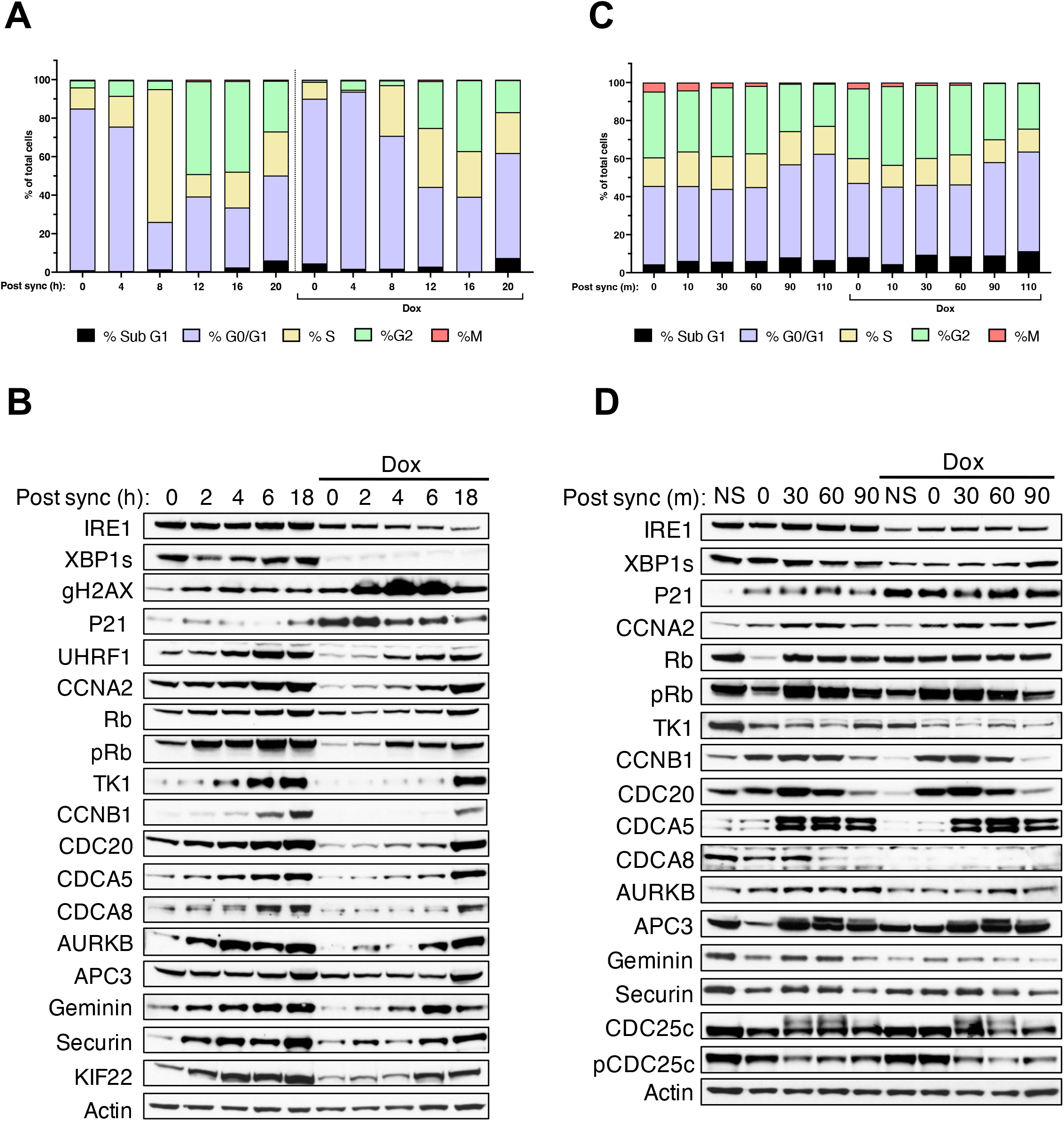
IRE1 depletion downregulates multiple cell cycle proteins. **(A)** Effect of IRE1 knockdown during G1 synchronization on cell cycle progression. AMO1 shIRE1 cl.1 cells were incubated with CDK4/6 inhibitor (1 μM) for 24 h in the absence or presence of Dox (0.2 μg /ml). Cells were washed to resume cycling in the absence or presence of Dox and at the indicated time points stained with PI and analyzed by flow cytometry for DNA content. Mitosis (M) was distinguished from G2 by phospho-H3 staining. **(B)** Effect of IRE1 knockdown during G1 synchronization on abundance of cell cycle proteins. AMO1 shIRE1 cl.1 cells were synchronized as in **A** in the absence or presence of Dox (0.2 μg /ml), washed to resume cycling in the absence or presence of Dox and analyzed at the indicated time points by IB. **(C)** Effect of IRE1 knockdown during late G2 synchronization on cell cycle progression. AMO1 shIRE1 cl.1 cells were incubated with CDK1 inhibitor (9 μM) for 20 h in the absence or presence of Dox (0.2 μg /ml). Cells were washed to resume cycling in the absence or presence of Dox and at the indicated time points stained with PI and analyzed by flow cytometry for DNA content. Mitosis (M) was distinguished from G2 by phospho-H3 staining. **(D)** Effect of IRE1 knockdown during late G2 synchronization on abundance of cell cycle proteins. AMO1 shIRE1 cl.1 cells were synchronized as in **C** in the absence or presence of Dox (0.2 μg /ml), washed to resume cycling in the absence or presence of Dox and analyzed at the indicated time points by IB.

Next, we synchronized AMO1 cells in late G2 by treatment with an established CDK1 inhibitor^66^, followed by inhibitor washout to allow mitotic progression. Flow cytometry indicated approximately 35% of the cells in G2 and another 5% in M phase at t=0 (**Fig. 7C and S7D**), totaling 40% as compared to the 20% of cells found in G2/M in non-synchronized cultures (**Fig. 2B**). Of note, the synchronization in G2/M was associated with somewhat increased levels of apoptosis overall. By 90 min post washout the proportion of cells in G1 began to increase, indicating completion of mitosis and re-entry into G1. IB analysis during the first 90 min after CDK1 inhibitor washout indicated a lower level of CDC20, CDCA8, AURKB, Geminin, and pCDC25c in IRE1-depleted cells vs. controls (**Fig. 7D**). Taken together, these results suggest that IRE1 coordinates both early and late events during cell cycle progression.

### Endogenous IRE1 protein can localize to the nuclear envelope

We reasoned that the regulation of cell cycle events by IRE1 might be facilitated by its localization to the nuclear envelope (NE), which has an outer membrane contiguous with ER membranes^67^. Supporting this hypothesis, subcellular fractionation of AMO1 cell lysates followed by IB revealed detectable IRE1 protein in the nuclear compartment, which contained the nuclear protein Lamin B1 but lacked the Golgi marker 58K (**Fig. 8A**). IRE1 knockdown depleted the protein in the nuclear fraction to a level below detection (**Fig. S8A**), confirming specificity. Furthermore, in contrast to Lamin B1, proteins containing the ER-localized epitope KDEL, and the ER-resident protein REEP5 were not detectable in the nuclear fraction, excluding the possibility that IRE1 detection in this compartment is due to ER contamination. To further investigate IRE1 localization, we leveraged a previously validated U2OS cell line expressing an endogenous IRE1 allele marked by CRISPR/CAS9-mediated gene editing with a C-terminal HaloTag^8^. Immuno-fluorescence microscopy using anti-HaloTag antibody suggested that, although most of the IRE1-HaloTag signal appeared to be associated with the ER, IRE1 protein could be detected in some proximity to Lamin B1, which marks the nuclear-matrix on the inner side of the NE (**Fig. 8B**). This proximity was further supported by integrated Z-stack analysis of 30 sections, each 300 nm thick (inset). To obtain more precise localization, we developed a sensitive immuno-EM technique to detect endogenous IRE1 similarly gene-edited with a HaloTag in AMO1 cells harboring Dox-inducible IRE1 shRNA. IB analysis of the modified AMO1 cells confirmed complete labeling of the endogenous IRE1 protein by the HaloTag, and its depletion upon Dox addition (**Fig. S8B**). Immuno-EM staining with the anti-HaloTag antibody and protein A Gold^10^ produced a relatively sparse signal; nevertheless, gold particles could be detected not only in association with ER membranes, but also with the NE (**Fig. 8C and Fig. S8C**). Thus, some of the cellular IRE1 protein can reside in the NE.

**Figure 8.**
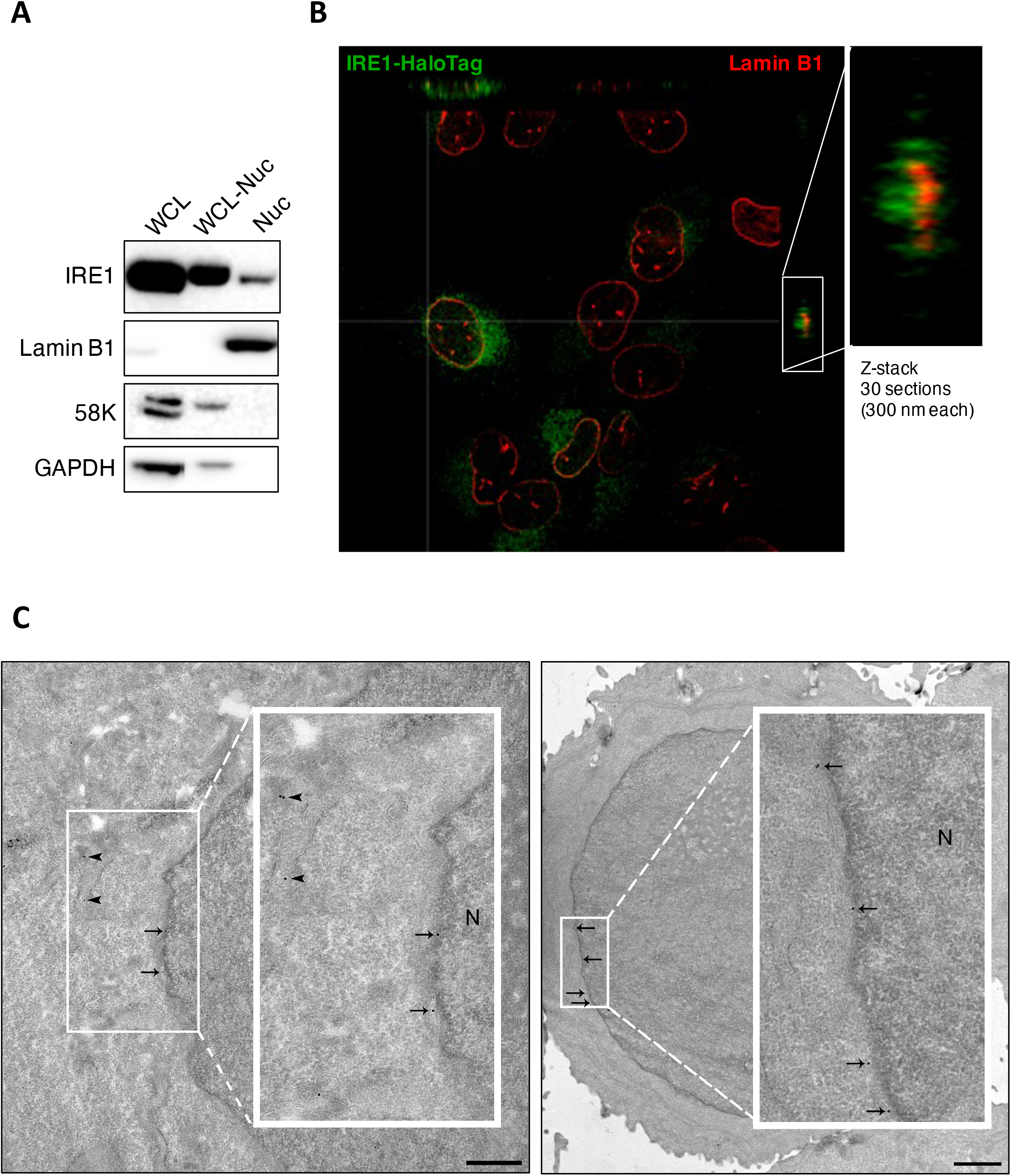
Endogenous IRE1 protein can localize to the nuclear envelope. **(A)** AMO1 cells were subjected to nuclear fractionation and analyzed by IB. WCL, whole-cell lysate. WCL-Nuc: Whole-cell lysate lacking the nuclear fraction. Nuc, nuclear fraction. The nuclear protein Lamin B1 and the Golgi marker 58K were used to confirm the purity of the nuclear fraction. **(B)** Detection of endogenous Halo-tagged IRE1 and non-tagged Lamin B1. U2OS HaloTag cells, containing a C-terminal HaloTag into the endogenous IRE1 gene locus^8^, were used. Cells were fixed, stained with anti-HaloTag (green) and anti-Lamin-B1 (red) antibodies and analyzed by confocal microscopy. Inset: Z-stack of 30 sections depicting colocalization between HaloTag and LaminB1. **(C)** Immuno-EM analysis of Halo-tagged endogenous IRE1 in AMO1 cells. A C-terminal HaloTag was inserted into the endogenous IRE1 gene locus in AMO1 cells as previously described^8^. Cells were fixed, stained with anti-HaloTag and protein-A-Gold10 nm conjugate (protein-A-Gold^10^), and analyzed by EM. Arrowheads and arrows indicate gold particles detected in association with ER membranes or nuclear envelope membranes, respectively. N = nucleus.

## DISCUSSION

Our results have potentially important implications for the current understanding of IRE1’s biological roles, as well as for the development of more effective strategies to disrupt IRE1 as a therapeutic target.

The evidence to date has suggested that cancer-cell dependency on IRE1 principally involves IRE1’s enzymatic function in the activation of XBP1s, with some possible additional contribution of RIDD^23,25,26^. Our present results demonstrate that some cancer cells can display a surprising nonenzymatic dependency on IRE1, without a significant requirement for XBP1s or RIDD. For certain tumor models, namely AMO1 and KMS27, IRE1 dependency was particularly strong, as evident by robust growth inhibition *in vitro* and complete tumor regression *in vivo* upon IRE1 silencing. We observed a nonenzymatic IRE1 requirement in both MM and colorectal carcinoma cells, suggesting that this phenotype is not restricted to one type of cancer. Aneuploidy and chromosomal instability are frequent hallmarks of malignancy^68^. Accordingly, it is conceivable that certain genome abnormalities, for example chromosomal translocations or other alterations, give rise to a unique dependency on IRE1 and a corresponding vulnerability to its disruption. The nature of such genomic alterations remains to be investigated.

Regardless, our data suggest that an as yet unidentified scaffolding role of IRE1—perhaps mediated by one or more of its known protein-protein associations, or through as yet unknown interactions—helps coordinate cell cycle progression. The ER plays important roles during the cell cycle, for example during mitotic breakdown and reassembly of the nuclear envelope^67^. Strikingly, our immuno-EM study of the endogenous IRE1 protein—to our knowledge the first such visualization—revealed that some IRE1 molecules can associate with the NE. Future studies should interrogate whether IRE1 fulfills a specific structural or scaffolding role in that location to facilitate ER-to-nucleus coordination during cell cycle progression. Intriguingly, previous IP-MS studies already suggested that IRE1 may directly interact with proteins involved in cell cycle regulation^22^.

IRE1 silencing led to a G1 cell cycle arrest as well as to apoptotic caspase activation, the former occurring independently of the latter. IRE1 depletion augmented DNA damage and chromosome instability, evident by an increased abundance of specific DDR markers and of micronuclei and a subsequent loss of DNA. These events may engage the p53 pathway to inhibit cell cycle progression across the G1-S as well as the G2-M checkpoints. CDKN1A/p21, found to be upregulated both at the mRNA and protein levels on IRE1 silencing, is a major transcriptional target of p53 that mediates cell cycle inhibition^57,58^. The p53 pathway also may contribute to the observed induction of apoptosis^69^. Indeed, proapoptotic transcriptional targets of p53 such as BAX and DR5/TNFRSF10B^70,71^ were upregulated upon knockdown of IRE1 but not XBP1 (**Fig. S4A**). Furthermore, the non-transcriptional upregulation of CDKN1B/p27 may provide additional cell cycle inhibition. A distinct interaction between the DDR and IRE1 has previously been reported based on evidence that DNA damage augments IRE1 activation and RIDD^72^. Furthermore, BRCA1—an E3 ubiquitin ligase involved in DNA repair—has been reported to promote proteostasis by ubiquitinating IRE1^73^. Together, these findings suggest a reciprocal connection between IRE1 and the DDR: while DNA damage enzymatically activates IRE1, IRE1 nonenzymatically promotes DNA integrity.

Another notable outcome of IRE1 silencing was the downregulation of UHRF1— a protein that acts as a key epigenetic regulator by bridging DNA methylation and chromatin modification^74^. UHRF1 specifically recognizes and binds hemimethylated DNA at replication forks and recruits the methyltransferase DNMT1 to ensure faithful propagation of DNA methylation patterns during S phase. UHRF1 also plays a key role in chromatin modification, binding to H3K9me3 and recruiting several chromatin-associated proteins^74,75^. IRE1 silencing decreased DNA methylation and attenuated 5-aza-induced H3K9me3 modification as well as heterochromatin abundance. Whether and how these phenotypes are connected to the downregulation of UHRF1 remains to be investigated.

Our data suggests that IRE1 fulfills a nonenzymatic structural function to support cell cycle progression, which is critical in certain cancer cells. More specifically, our transcriptomic and proteomic analyses indicate that IRE1 regulates cell cycle events that promote DNA replication, chromosome segregation, as well as chromatin maintenance. IRE1 silencing downregulated mRNA transcripts and/or proteins involved in advancing S phase as well as mitosis. It is possible that the inhibition of S-phase by IRE1 depletion causes mitotic disruption within the same cell cycle. Alternatively, perturbations of DNA integrity may lead to G1 arrest of afflicted daughter cells during the subsequent cycle^76^. Perhaps related to our present findings is published evidence that IRE1 supports mitotic fidelity of chromosome segregation in the budding yeast *S. cerevisiae*^77^. Further data indicates the existence of an ER stress surveillance mechanism that ensures proper ER inheritance during cell division in budding yeast^78,79^. In addition, IRE1 has been implicated in regulating sensitivity of KRAS-mutant colon cancer cells to MEK inhibition^40^, and in resistance of pancreatic cancer cells to direct KRAS-mutant inhibition^43^. Additional evidence with potential relevance to the present results include the translocation of proteolytic IRE1 fragments from the ER to the nucleus through cleavage by Presenilin-1^80^, and the genetic demonstration of a critical nonezymatic IRE1 requirement for VDJ recombination and B-cell receptor generation in mouse pre-B cells^81^.

Perhaps the most significant translational implication of this study is that enzymatic IRE1 inhibition does not suffice to realize IRE1’s full potential as a therapeutic target for cancer. Earlier work has shown that enzymatic inhibition of IRE1, either at the kinase or the RNase level, or XBP1 disruption, can be efficacious in models representing diverse cancers. Herein we reveal that enzymatic IRE1 inhibition or XBP1 knockdown fails to restrain malignant growth of certain cancer models that are nevertheless attenuated by IRE1 silencing. This generates a strong conceptual rationale to explore novel disruptive strategies for IRE1 at the gene, mRNA, or protein level, *e.g.*, gene therapy, antisense oligonucleotides, small-interfering RNAs, or protein degraders. Such interventions may be more broadly effective than IRE1 kinase or RNase inhibition and should be considered for cancer and perhaps for other pathologies that may involve nonenzymatic IRE1 function. Furthermore, specific biomarker strategies are needed to better predict IRE1 dependency in order to guide optimal IRE1-disruptive therapy for the individual patient.

## FIGURE LEGENDS

**Figure S1.**
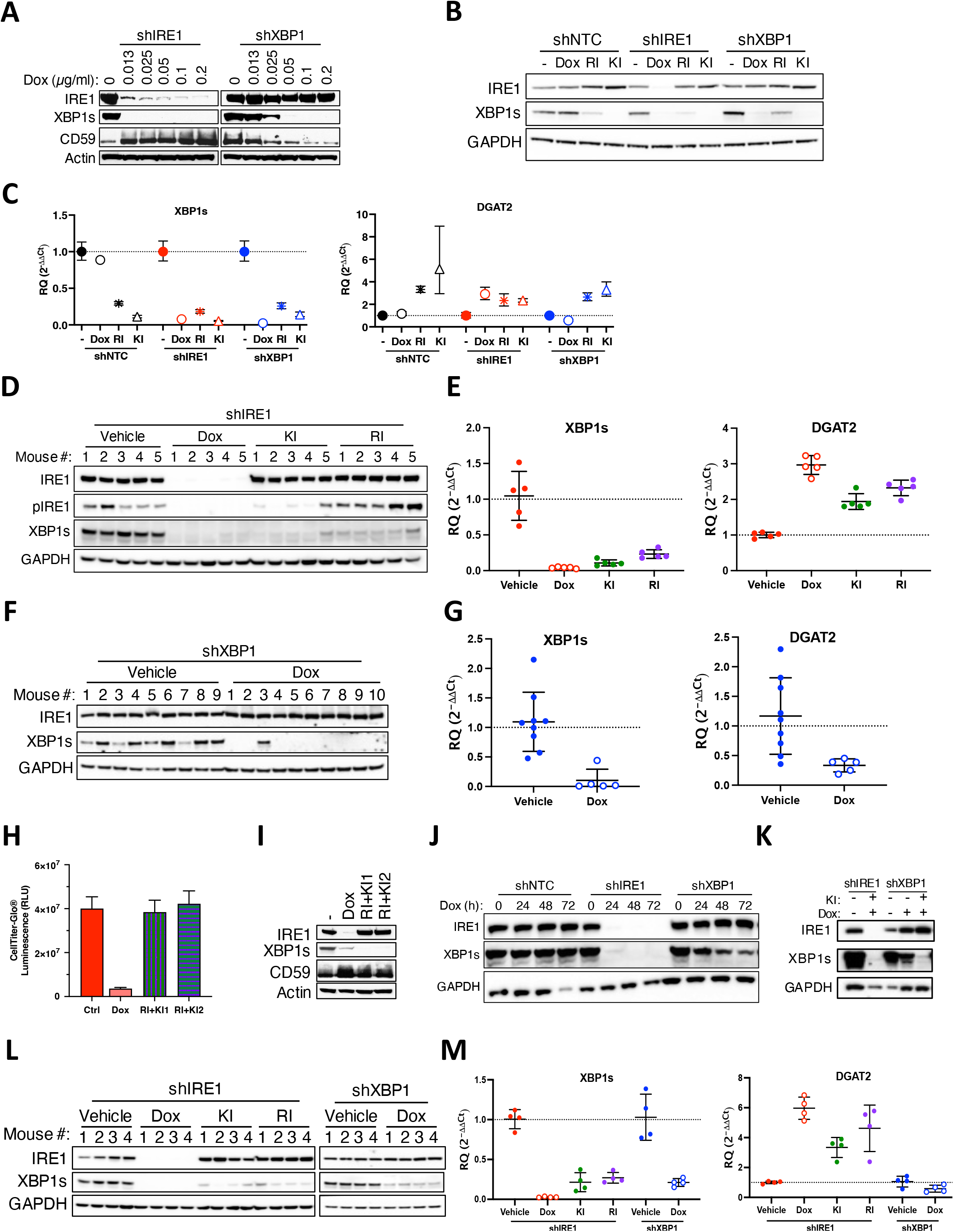
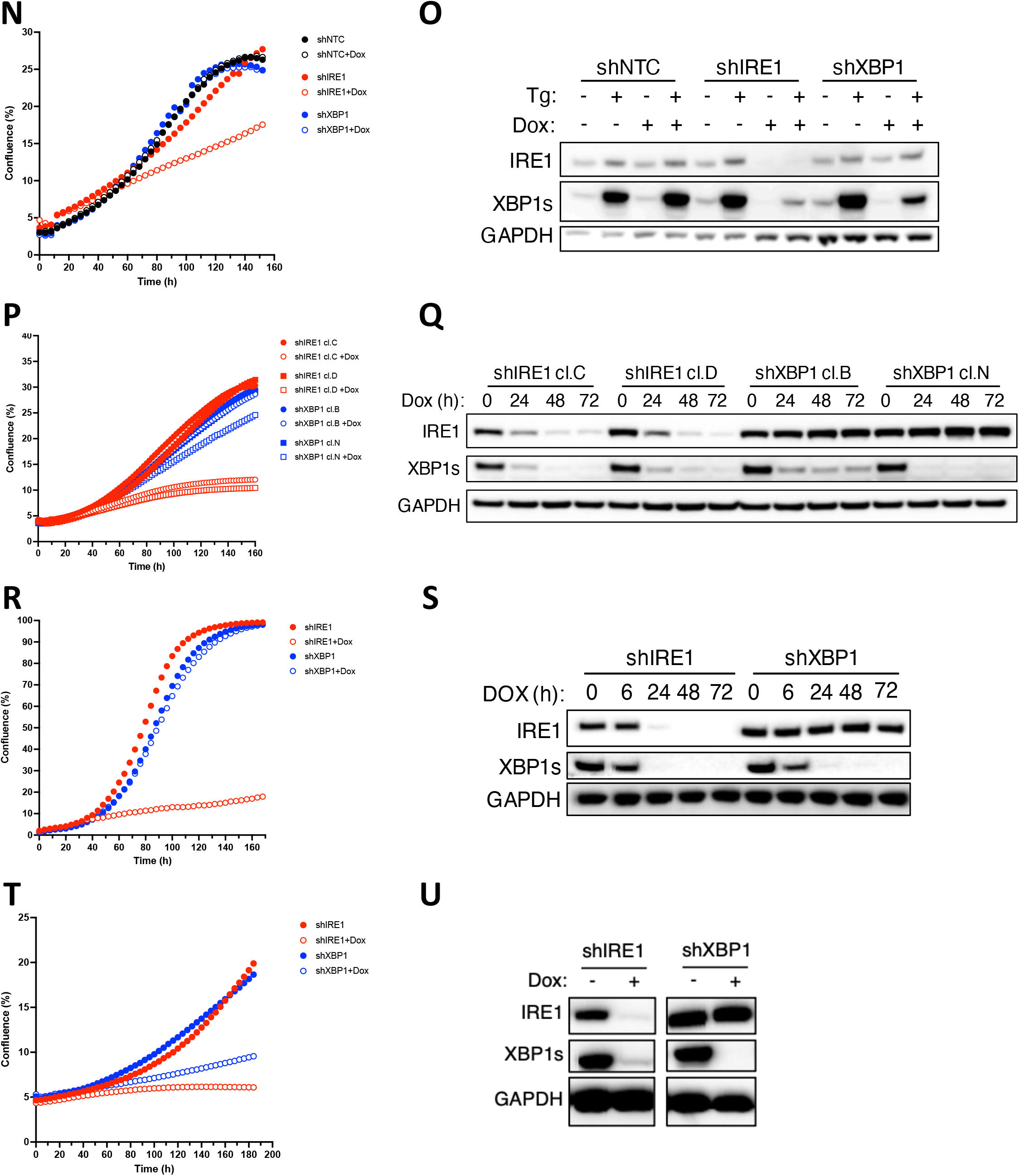
Some cancer cell lines require IRE1 but not its enzymatic activity. **(A)** Validation of IRE1 and XBP1s knockdown in AMO1 cells *in vitro*. Cells stably transfected with plasmids encoding Dox-inducible shRNAs against IRE1 or XBP1 were incubated for 48 h in the absence or presence of Dox at the indicated concentrations. Cells were then analyzed by Immunoblotting (IB). One independent clone for each gene (cl.1 for shIRE1 and cl.1 for shXBP1) is shown. **(B)** Validation of shRNA knockdown and enzymatic IRE1 inhibition in AMO1 cells. AMO1 shNTC, shIRE1 cl.1 and shXBP1 cl.1 cells were incubated for 48 h in the absence or presence of Dox (0.2 μg/ml), or IRE1 RI (3 μM), or IRE1 KI (KI2, 3 μM). Cells were analyzed by IB. Because XBP1s production requires both IRE1 kinase and RNase activity, its depletion confirms the inhibition of both functions. **(C)** Cells were treated as in **B** and analyzed by RT-qPCR for mRNA levels of *XBP1s* and *DGAT2* (RIDD target). **(D)** Validation of shRNA knockdown and enzymatic IRE1 inhibition in AMO1 cells *in vivo*. C.B-17 SCID mice were implanted subcutaneously with 10×10^6^ AMO1 shIRE1 cl.1 cells and allowed to form palpable tumors. Mice were then randomized into treatment groups (n=5/group) and given either vehicle (5% sucrose) or Dox (0.5 mg/ml in 5% sucrose) *ad libitum*, or treated orally bidaily with vehicle, or IRE1 KI (250 mg/kg) or IRE1 RI (100 mg/kg) over 3.5 days. 6h after the last oral dosing animals were sacrificed, tumors collected and protein extracted for IB analysis. **(E)** Mice were treated as in **D** and tumor samples were analyzed by RT-qPCR for XBP1s and DGAT2 mRNA levels as in **C**. **(F)** Validation of XBP1 knockdown *in vivo* in AMO1 tumor xenografts. Tumors were collected at endpoint from the study depicted in Fig. 1B. Tumor samples were analyzed by IB. **(G)** Tumor samples shown in **F** were analyzed as in **C** by RT-qPCR. **(H)** Effect of IRE1 knockdown or KI or RI or KI+RI treatment on *in vitro* growth of AMO1 cells. Cells were treated with Dox (0.2 μg/ml), or IRE1 KI1 (G5758) (1 μM) or KI2 (G9668) (1 μM), or IRE1 RI (1 μM), or both (1 μM each) for 72 h and analyzed by CellTiter-Glo. **(I)** IB validation of IRE1 depletion or enzymatic inhibition for panel **H**. **(J)** Effect of IRE1 or XBP1 knockdown in KMS27 cells *in vitro*. Cells stably transfected with plasmids encoding Dox-inducible shRNAs against NTC or IRE1 or XBP1 were incubated for the indicated time in the absence or presence of Dox (0.2 μg/ml) and analyzed by IB. **(K)** Validation of knockdown and enzymatic IRE1 inhibition *in vitro* in KMS27 cells. Cells were incubated in the absence or presence of 0.2 μg/ml Dox or IRE1 KI (KI2, 3 μM) for 72 h and analyzed by IB. **(L)** Validation of knockdown and enzymatic IRE1 inhibition *in vivo* in KMS27 tumors. C.B-17 SCID mice implanted subcutaneously with 10×10^6^ KMS27 shIRE1 cl.9 or shXBP1 cl.13 cells and allowed to form palpable tumors. Mice were then randomized into treatment groups (n=4/group) and given either vehicle (5% sucrose) or Dox (0.5 mg/ml in 5% sucrose) *ad libitum*, or treated orally bidaily with either vehicle, IRE1 KI (250 mg/kg) or IRE1 RI (100 mg/kg), over 3.5 days. 6h after the last oral dosing animals were sacrificed, tumors collected and protein extracted for IB analysis. **(M)** RT-qPCR analysis of XBP1s and DGAT2 mRNA in tumor samples described in **L**. **(N)** Effect of IRE1 or XBP1 knockdown on *in vitro* proliferation of JJN3 cells. Cells were stably transfected with a plasmid encoding Dox-inducible shRNAs against NTC (black) IRE1 (red) or XBP1 (blue). Cells were incubated in the absence (closed symbols) or presence (open symbols) of Dox (0.2 μg/ml) and analyzed for proliferation as in Fig. 1A. **(O)** Validation of IRE1 and XBP1 knockdown in JJN3 cells. Cells described in **N** were incubated with Dox (0.2 μg/ml) for 48 h and analyzed by IB. Thapsigargin (Tg) was added for 1h at 100nM to induce IRE1 activity. **(P)** Effect of IRE1 or XBP1 knockdown on *in vitro* proliferation of L363 cells. Cells were stably transfected with plasmids encoding Dox-inducible shRNAs against IRE1 (red) or XBP1 (blue). Two independent clones (circles or squares) for each gene knockdown were characterized. Cells were incubated in the absence or presence of Dox (0.2 μg/ml) and analyzed for proliferation as in Fig. 1A. **(Q)** Validation of knockdown in L363 cells. Cells described in **P** were incubated with Dox (0.2 μg/ml) for the indicated time and analyzed by IB. **(R)** Effect of IRE1 or XBP1 knockdown on *in vitro* proliferation of HCT116 cells. Cells were stably transfected with plasmids encoding Dox-inducible shRNAs against IRE1 (red) or XBP1 (blue). Cells were incubated in the absence (closed symbols) or presence (open symbols) of Dox (0.2 μg/ml) and analyzed for proliferation as in Fig. 1A. **(S)** Validation of knockdown in HCT116 cells. Cells described in **R** were incubated with Dox (0.2 μg/ml) for the indicated time and analyzed by IB. **(T)** Effect of IRE1 or XBP1 knockdown on *in vitro* proliferation of OPM2 cells. Cells were stably transfected with plasmids encoding Dox-inducible shRNAs against IRE1 or XBP1. Cells were incubated in the absence or presence of Dox (0.2 μg/ml) and analyzed for proliferation as in Fig. 1A. **(U)** Validation of knockdown in OPM2 cells. Cells described in **T** were incubated with Dox (0.2 μg/ml) for 72 h and analyzed by IB.

**Figure S2.**
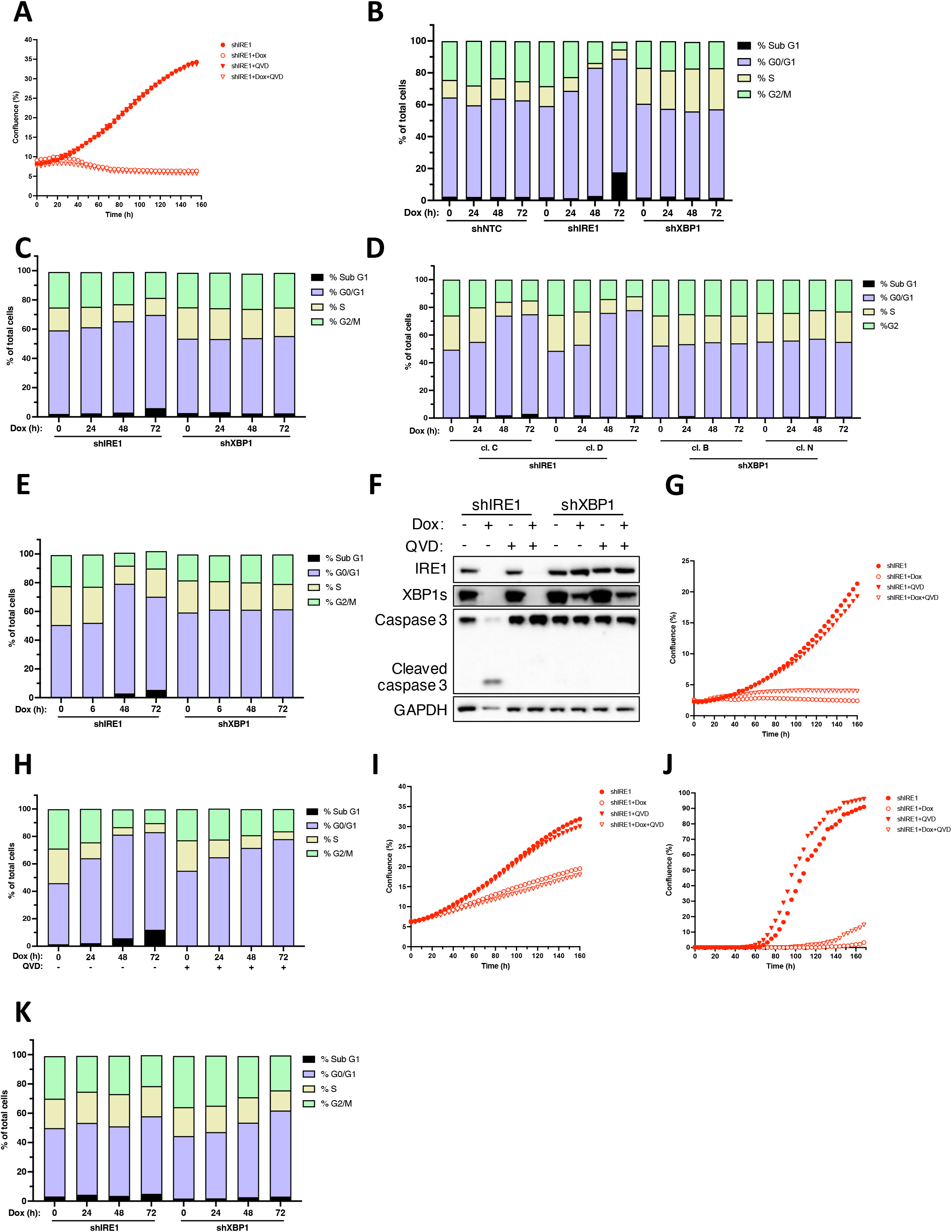
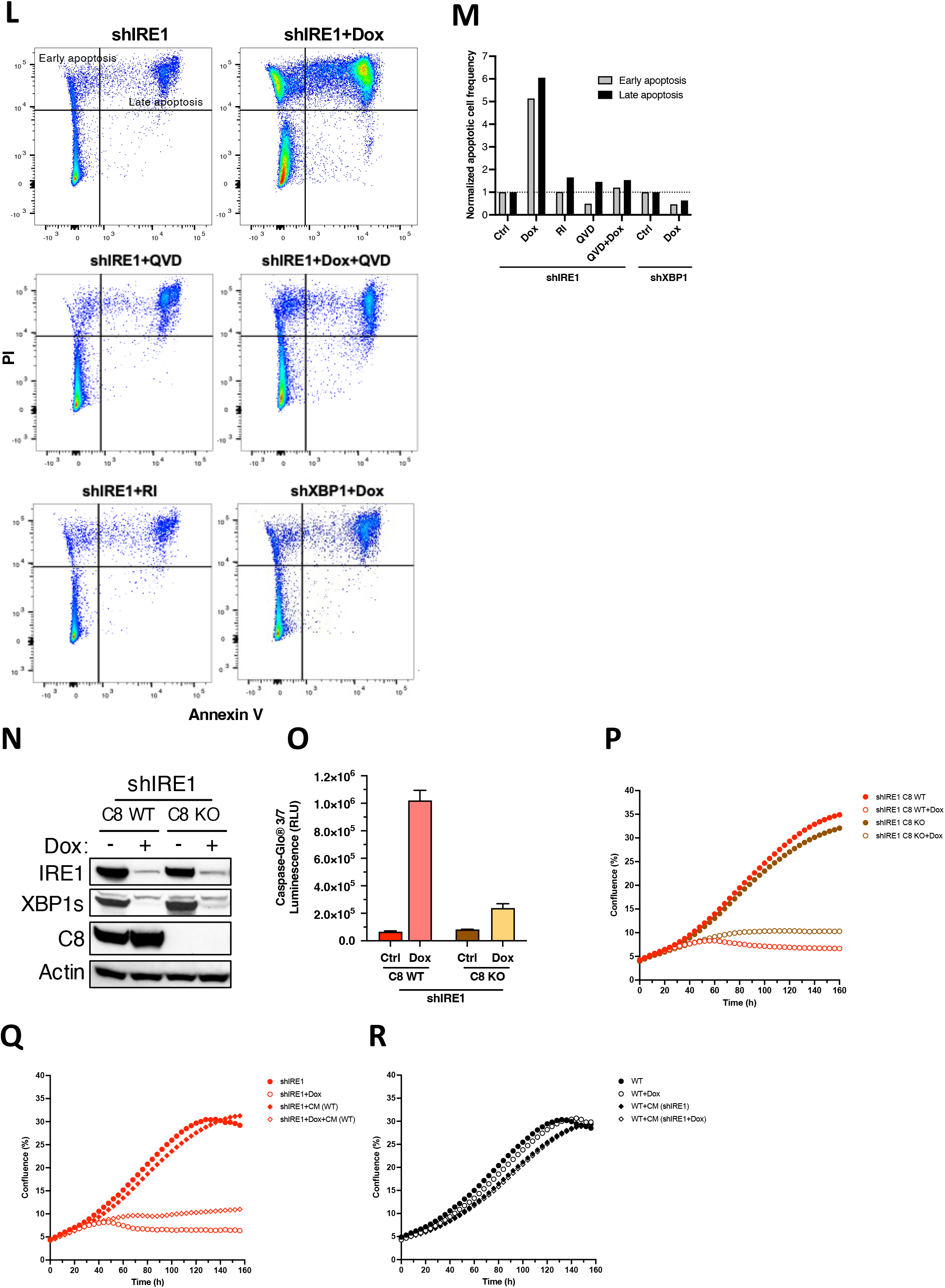
IRE1 depletion induces cell cycle arrest independently of apoptosis. **(A)** Effect of QVD on AMO1 cell proliferation. AMO1 shIRE1 cl.1 cells were treated for 24 h with Dox (open symbols) (0.2 μg/ml) in the absence (circles) or presence (triangles) of QVD (30 μM) and analyzed for proliferation. Proliferation, depicted as % confluence, was monitored by time-lapse microscopy in an Incucyte^TM^ instrument. **(B)** Effect of IRE1 or XBP1 knockdown on cell cycle progression. KMS27 shIRE1 cl.9 or shXBP1 cl.13 cells were incubated in the absence or presence of Dox (0.2 μg/ml) for the indicated time, stained with propidium iodide (PI) and analyzed by flow cytometry to determine cell cycle phase by DNA content. **(C)** JJN3 cells stably transfected with plasmids encoding Dox-inducible shRNAs against IRE1 or XBP1 were incubated with Dox (0.2 μg/ml) for the indicated time, stained with PI and analyzed by flow cytometry as in **B**. **(D)** L363 cells stably transfected with plasmids encoding Dox-inducible shRNAs against IRE1 (cl.C and cl.D) or XBP1 (cl. B and cl.N) were incubated with Dox (0.2 μg/ml) for the indicated time, stained with PI and analyzed by flow cytometry as in **B**. **(E)** HCT116 cells stably transfected with plasmids encoding Dox-inducible shRNAs against IRE1 or XBP1 were incubated for the indicated time with Dox (0.2 μg/ml), stained with PI and analyzed by flow cytometry as in **B**. **(F)** KMS27 shIRE1 cl.9 or shXBP1 cl.13 cells were incubated for 24 h with Dox (0.2 μg/ml) in the absence or presence of QVD (30 μM) and analyzed by IB. **(G)** KMS27 shIRE1 cl.9 or shXBP1 cl.13 cells were treated as in **F** and analyzed for confluence as in **A**. **(H)** KMS27 shIRE1 cl.9 or shXBP1 cl.13 cells were treated as in **F** for the indicated time, stained with PI and analyzed by flow cytometry as in **B**. **(I)** JJN3 shIRE1 cells were incubated with Dox (0.2 μg/ml) in the absence or presence of QVD (30 μM) and analyzed for confluence as in **A**. **(J)** HCT116 shIRE1 cells were incubated with Dox (0.2 μg/ml) in the absence or presence of QVD (30 μM) and analyzed for confluence as in **A**. **(K)** OPM2 cells stably transfected with plasmids encoding Dox-inducible shRNAs against IRE1 or XBP1 were incubated with Dox (0.2 μg/ml) for the indicated time, stained with PI and analyzed by flow cytometry as in **B**. **(L)** AMO1 shIRE1 cl.1 or shXBP1 cl.1 cells were incubated for 72 h with Dox (0.2 μg/ml) or IRE1 KI or RI (3 μM) in the absence or presence of QVD (30 μM), stained with PI and FITC anti-Annexin V antibody and analyzed by flow cytometry for early (low PI, high FITC anti-Annexin V) or late apoptosis (high PI and FITC anti-Annexin V), as indicated in the quadrants. **(M)** Graph depicting the frequency of cells shown in **L** in early apoptosis and late apoptosis. Results shown as fold change of apoptotic cells to the untreated controls. **(N)** Validation of caspase-8 knockout in AMO1 cells. AMO1 shIRE1 cl.1 cells were subjected to CRISPR/CAS9-mediated disruption using guide RNA against caspase-8 (C8). Parental (C8 WT) or C8 knockout (C8 KO) cells were incubated for 72 h in the absence or presence of Dox (0.2 μg/ml) and analyzed by IB. **(O)** AMO1 shIRE1 cl.1 C8 WT or C8 KO cells were incubated for 72 h with Dox (0.2 μg/ml) and analyzed for caspase activity by caspase-3/7 Glo assay. **(P)** AMO1 shIRE1 cl.1 C8 WT or C8 KO cells were incubated with Dox (0.2 μg/ml) and analyzed for confluence as in **A**. **(Q)** Effect of conditioned media on growth of AMO1 cells. Conditioned media were collected after 24 h of culture from naïve AMO1 cells [CM (WT)]. AMO1 shIRE1 cl.1 cells were then grown in the absence or presence of Dox (0.2 μg /ml) together with fresh media or CM (WT) and analyzed for confluence as in **A**. **(R)** CM were collected after 24 h of culture from AMO1 shIRE1 cl.1 cells grown in the absence or presence of Dox (0.2 μg /ml) [(CM (shIRE1) or CM (shIRE1+Dox), respectively]. AMO1 WT cells were then grown in the absence or presence of Dox, or CM (shIRE1) or CM (shIRE1+Dox) and analyzed for confluence as in **A**.

**Figure S3.**
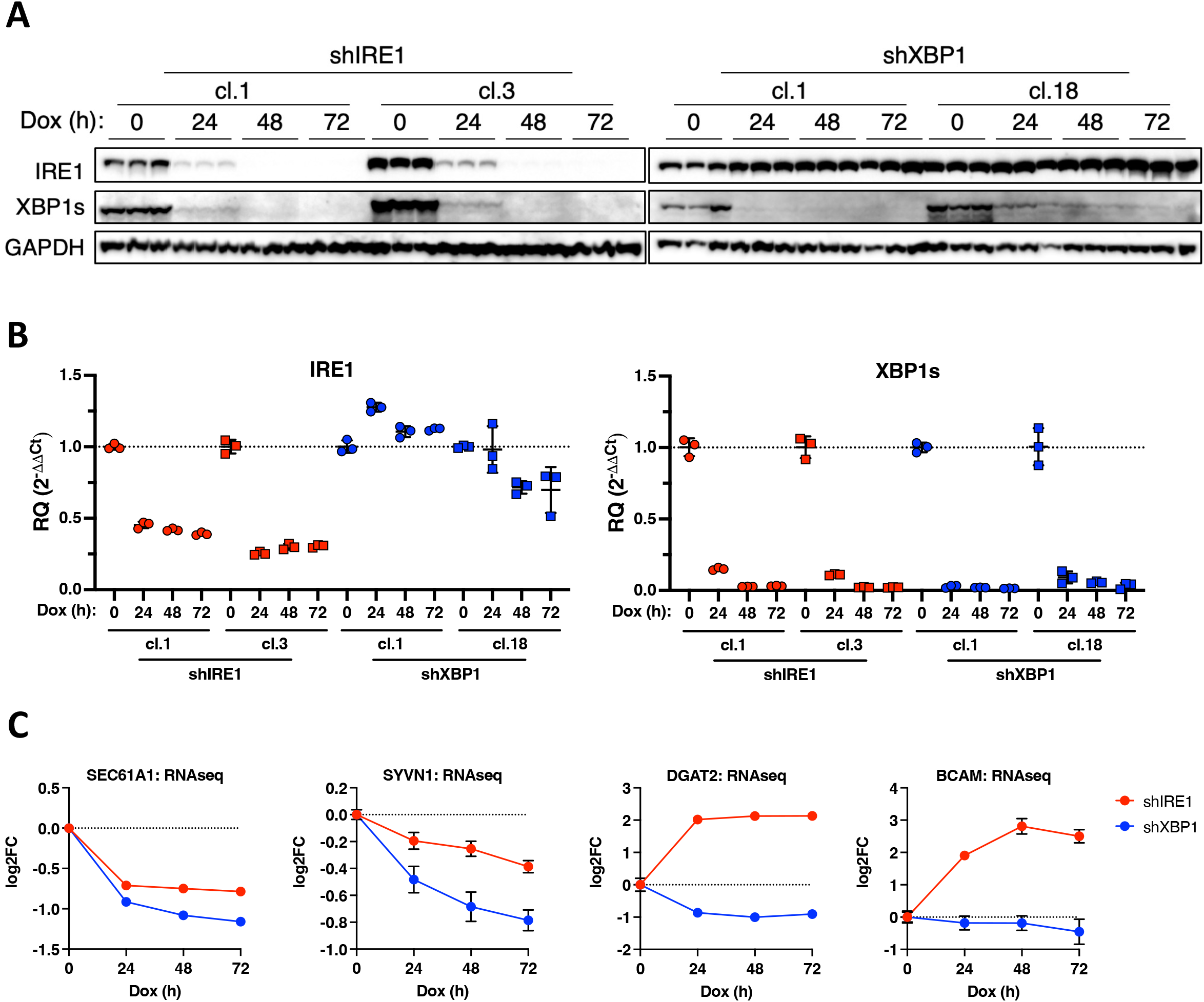
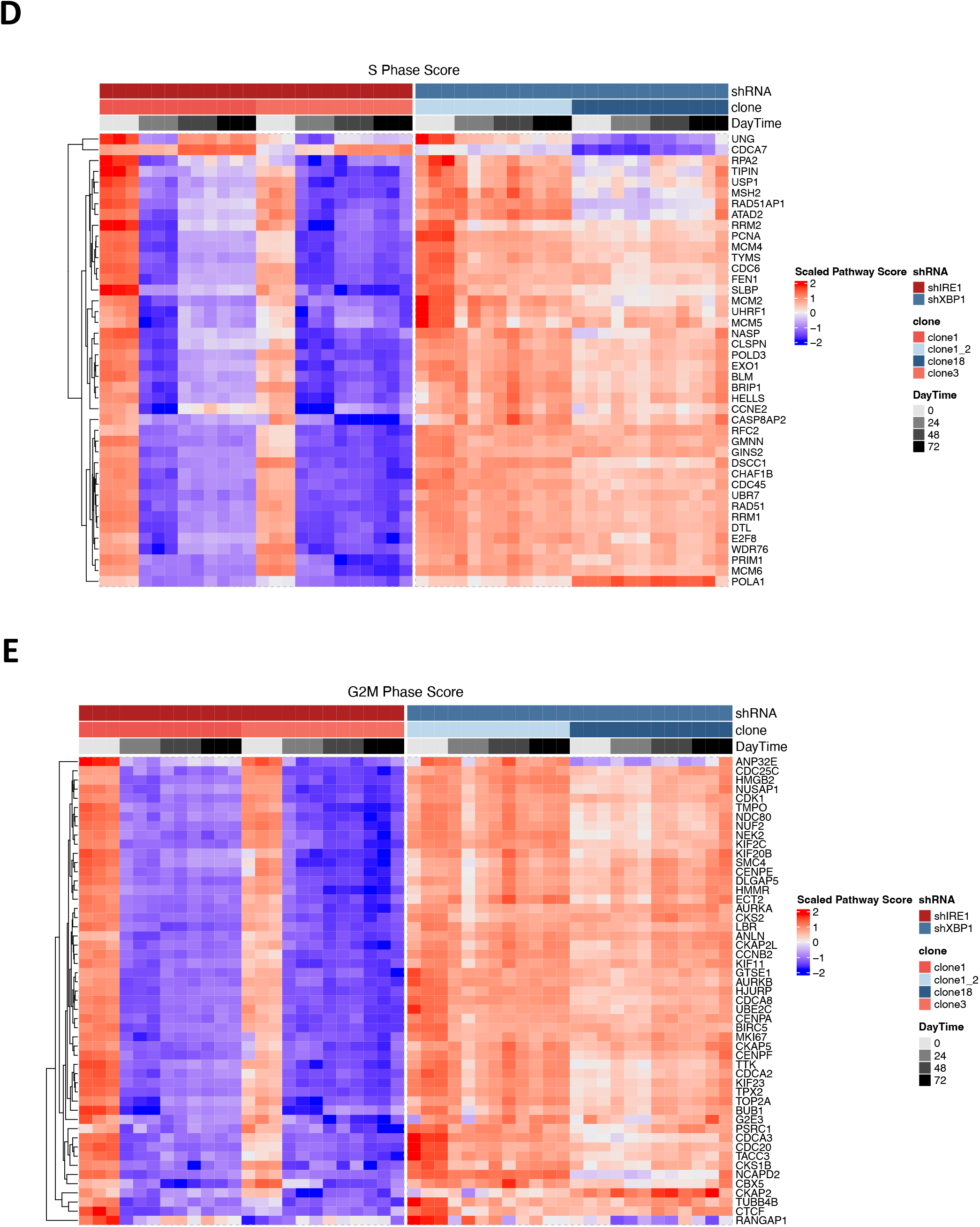
IRE1 silencing downregulates expression of multiple cell cycle genes. **(A)** RNA sequencing sample validation. AMO1 shIRE1 cl.1 or cl.3 cells and shXBP1 cl.1 and cl.18 cells were incubated for the indicated time with Dox (0.2 μg /ml) in triplicates and analyzed by IB. **(B)** Cells as in **A** were analyzed by RT-qPCR. **(C)** Effect of IRE1 or XBP1 knockdown on mRNA expression of select XBP1s- and RIDD-target genes. Cells as in **A** were analyzed by bulk RNA sequencing (RNAseq) for mRNA expression of the XBP1s targets *Sec61A1* and *SYVN1* or the RIDD targets *DGAT2* and *BCAM*. **(D)** Heatmap of the mRNA expression by bulk RNAseq of genes involved in the S phase of the cell cycle from RNA sequencing of cells as in **A**, used to calculate the S phase score in Fig. 3D. **(E)** Heatmap of the mRNA expression by bulk RNAseq of genes involved in the G2 and M phases of the cell cycle from RNA sequencing of cells as in **A**, used to calculate the G2/M phase score in Fig. 3D.

**Figure S4.**
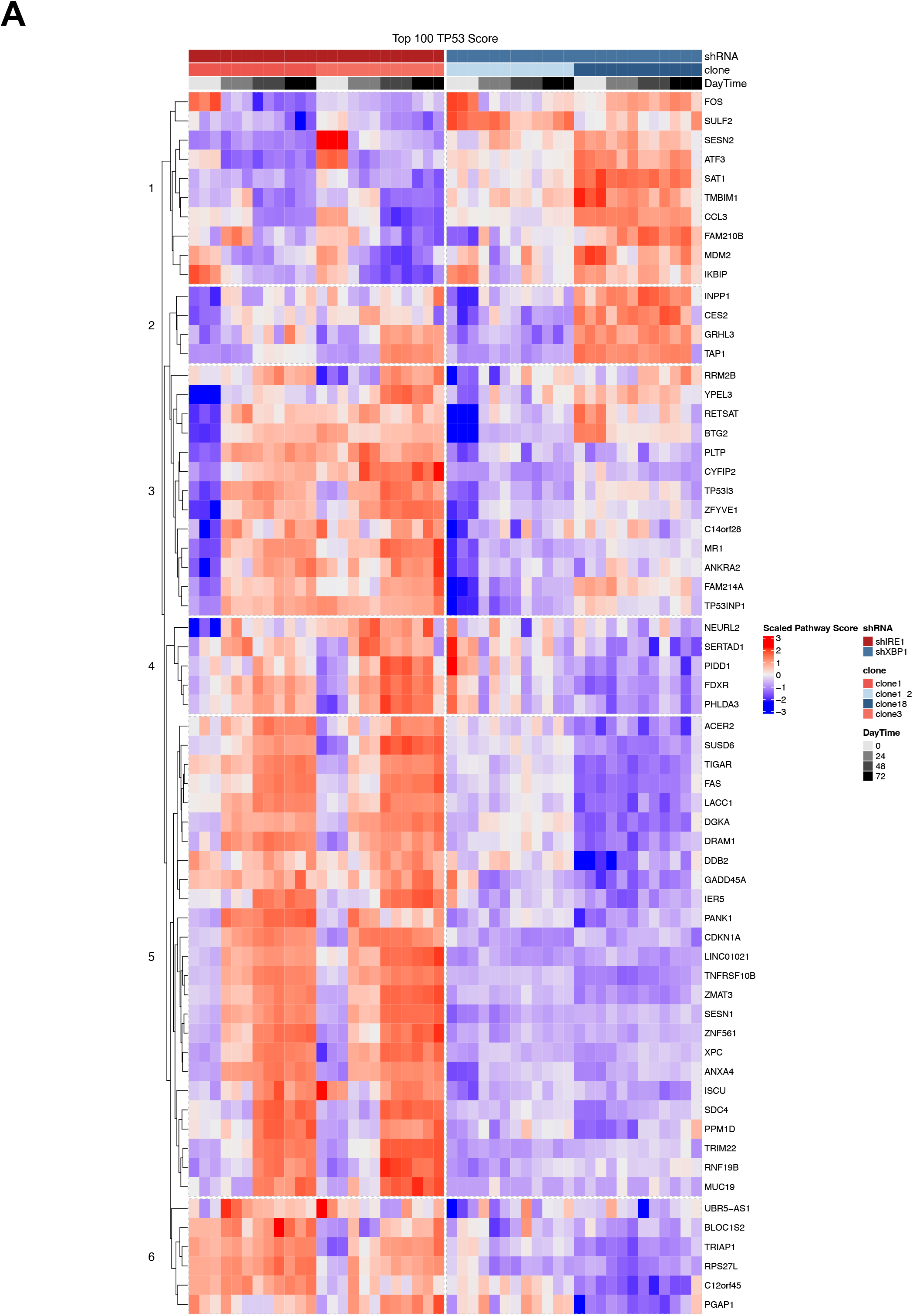
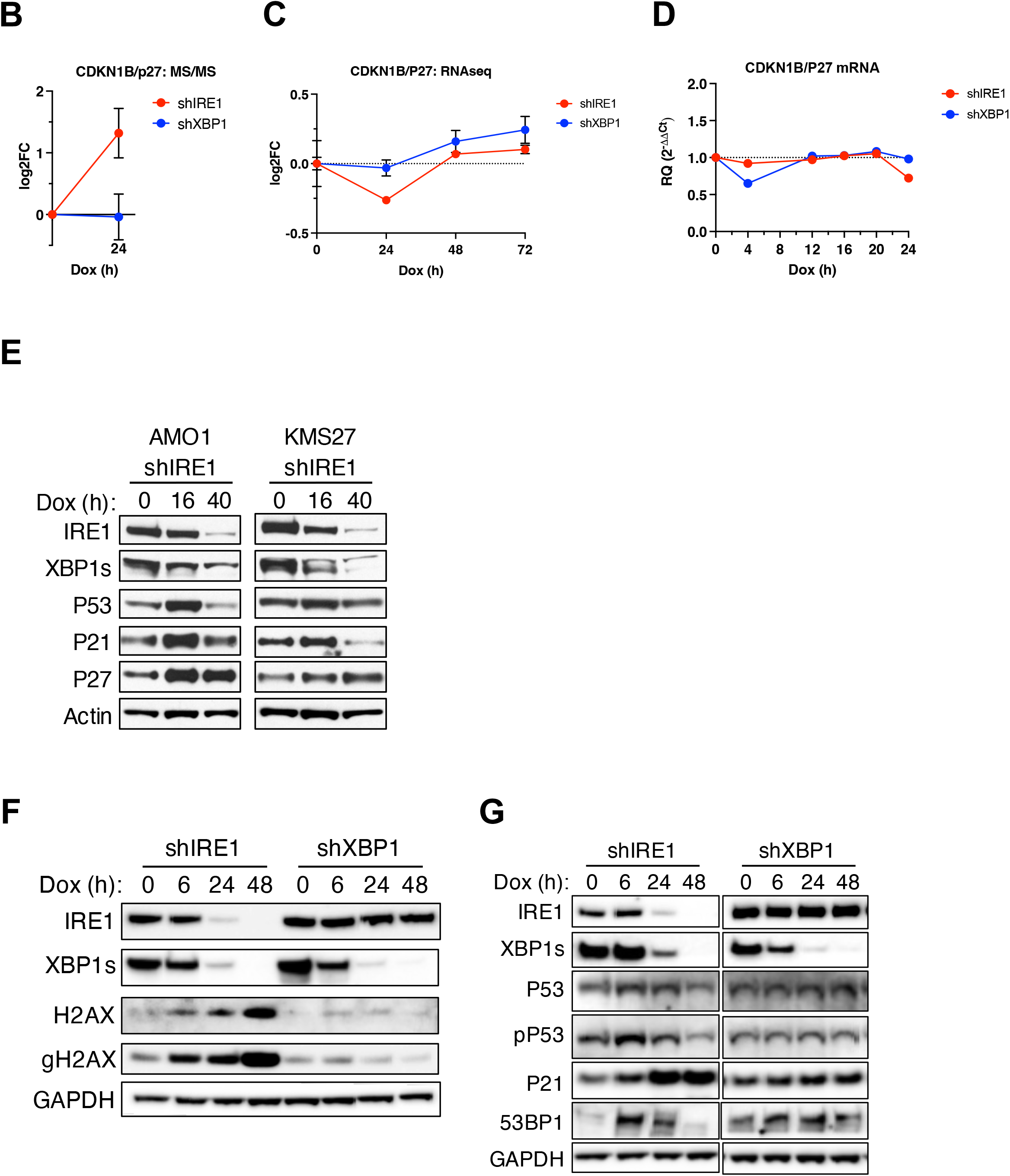
IRE1 depletion upregulates the p53 pathway and specific CDK inhibitors. **(A)** Effect of IRE1 or XBP1 knockdown on mRNA expression of TP53 target genes. Heatmap depicting mRNA expression by RNA sequencing of the top 100 TP53 pathway genes for samples depicted in Fig. 3A. **(B)** Effect of IRE1 or XBP1 knockdown on CDKN1B/p27 protein levels. Data depicted are from the proteomics analysis described in Fig. 4B. **(C)** Effect of IRE1 or XBP1 knockdown on CDKN1B/p27 mRNA levels. Data depicted are from RNAseq analysis described in Fig. 3A. **(D)** Effect of IRE1 or XBP1 knockdown on p27 mRNA levels. AMO1 shIRE1 cl.1 or shXBP1 cl.1 cells were incubated for the indicated time with Dox (0.2 μg /ml) and analyzed by RT-qPCR. **(E)** Effect of IRE1 knockdown on p53, p21 and p27 protein levels. AMO1 shIRE1 cl.1 and KMS27 shIRE1 cl.9 cells were incubated for the indicated time with Dox (0.2 μg /ml) and analyzed by IB. **(F)** Effect of IRE1 or XBP1 knockdown on H2AX and phosphorylated (γ) H2AX (gH2AX). AMO1 shIRE1 cl.1 or shXBP1 cl.1 cells were incubated for the indicated time with Dox (0.2 μg /ml) and analyzed by IB. **(G)** Effect of IRE1 or XBP1 knockdown on p53, p21 and 53BP1 proteins. AMO1 shIRE1 cl.1 or shXBP1 cl.1 cells were incubated for the indicated time with Dox (0.2 μg /ml) and analyzed by IB.

**Figure S5.**
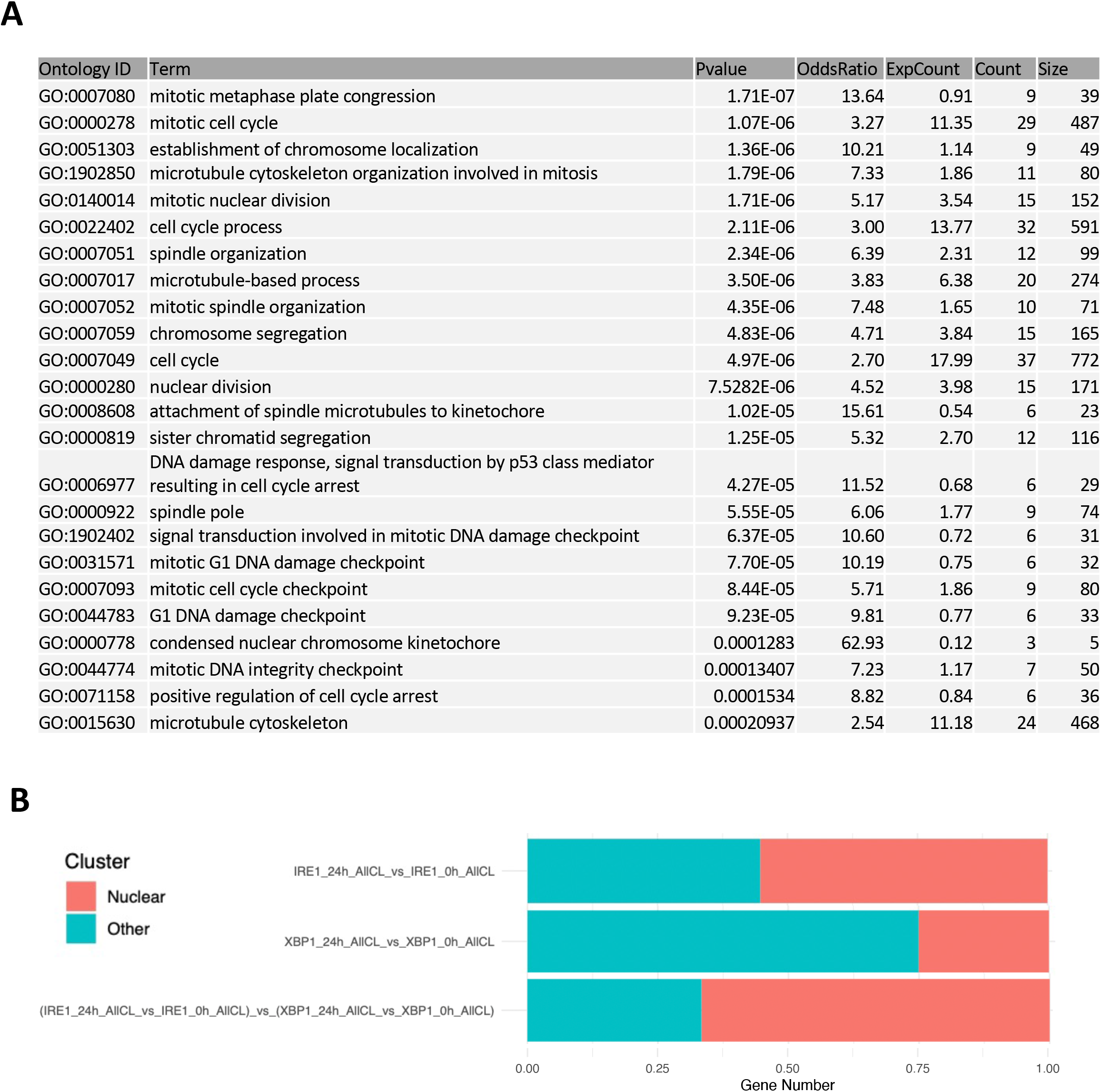
IRE1 increases chromosome instability. **(A)** Significant pathways from the Gene Ontology (GO) analysis of protein expression from the proteomics experiment shown in Fig. 4B. **(B)** Effect of IRE1 or XBP1 knockdown on the fraction of nuclear vs non-nuclear localizing proteins based on proteomics results from Fig. 4B.

**Figure S6.**
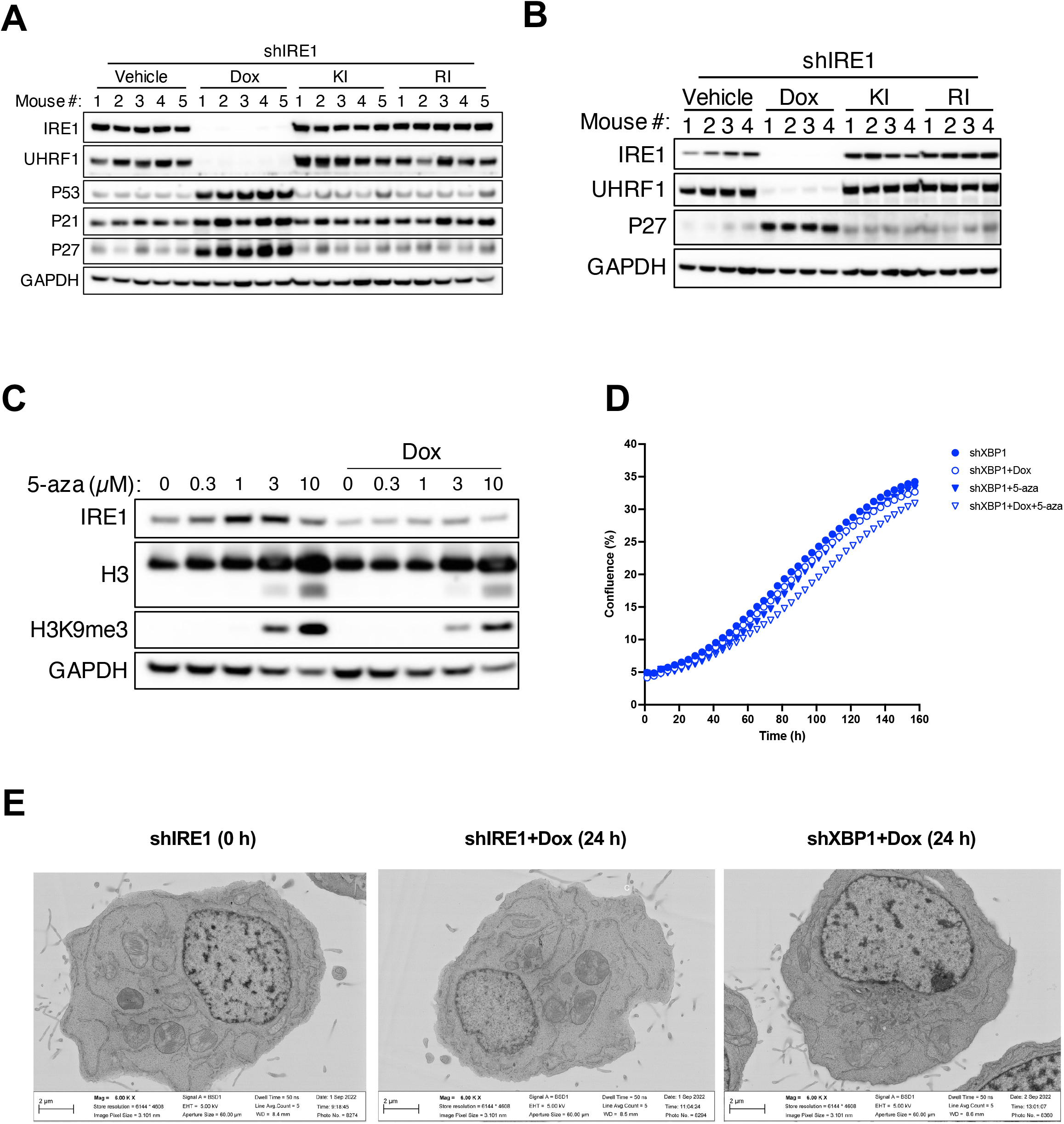
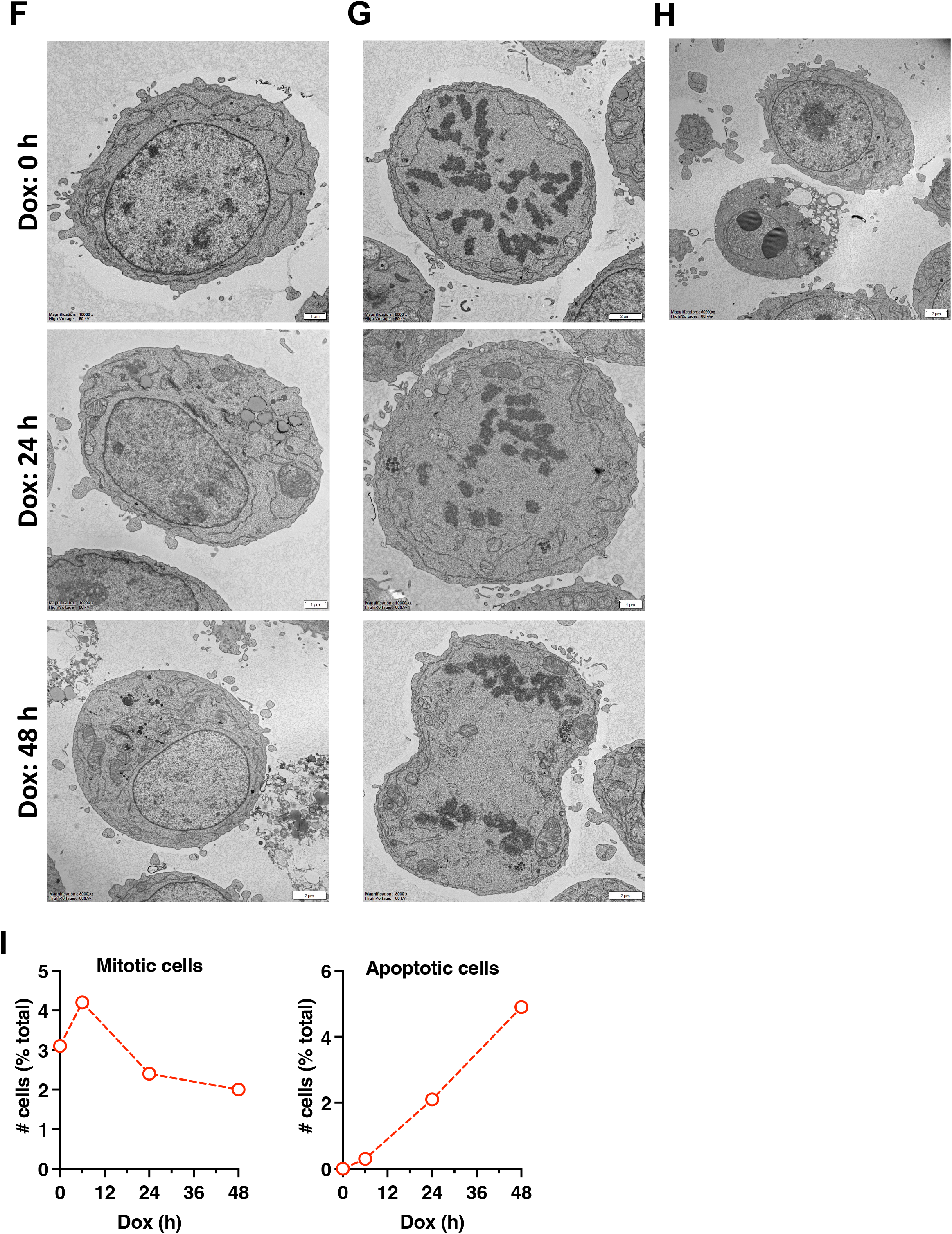
IRE1 silencing decreases UHRF1, DNA and H3 methylation and heterochromatin. **(A)** Effect of IRE1 or XBP1 knockdown *in vivo* on UHRF1 protein levels. AMO1 tumor xenografts as depicted in **Fig. S1D** were analyzed by IB. The IRE1 and GAPDH blots from **Fig. S1D** are duplicated here. **(B)** KMS27 tumor xenografts as depicted in **Fig. S1L** were analyzed by IB. The IRE1 and GAPDH blots from **Fig. S1L** are duplicated here. **(C)** AMO1 shIRE1 cl.1 cells were incubated for 24 h in the absence or presence of Dox (0.2 μg /ml) with the indicated concentration of 5-aza and analyzed by IB. The quantification for these IBs is depicted in Fig. 6D. **(D)** Effect of XBP1 knockdown and 5-aza treatment on proliferation. AMO1 shXBP1 cl.1 cells were incubated and analyzed as in Fig. 6E. **(E)** Additional examples of cells analyzed as depicted in **Fig. 6F and G**. **(F)** Examples of cells analyzed by EM (Utrecht) for heterochromatin quantification as depicted in **Fig. 6F** and **G** and used for the quantification in Fig. 6H. Scale bars are 1 μm. **(G)** Examples of mitotic cells analyzed by EM (Utrecht) for the quantification shown in **I**. Scale bars are 1 μm. **(H)** Example of an apoptotic cell analyzed by EM (Utrecht) for quantification shown in **I**. Scale bars are 1 μm. **(I)** Frequency of mitotic and apoptotic cells analyzed by EM (Utrecht) in AMO1 shIRE1 cl.1 cells incubated for the indicated time with Dox (0.2 μg /ml). For each sample 164 - 196 cells were examined.

**Figure S7.**
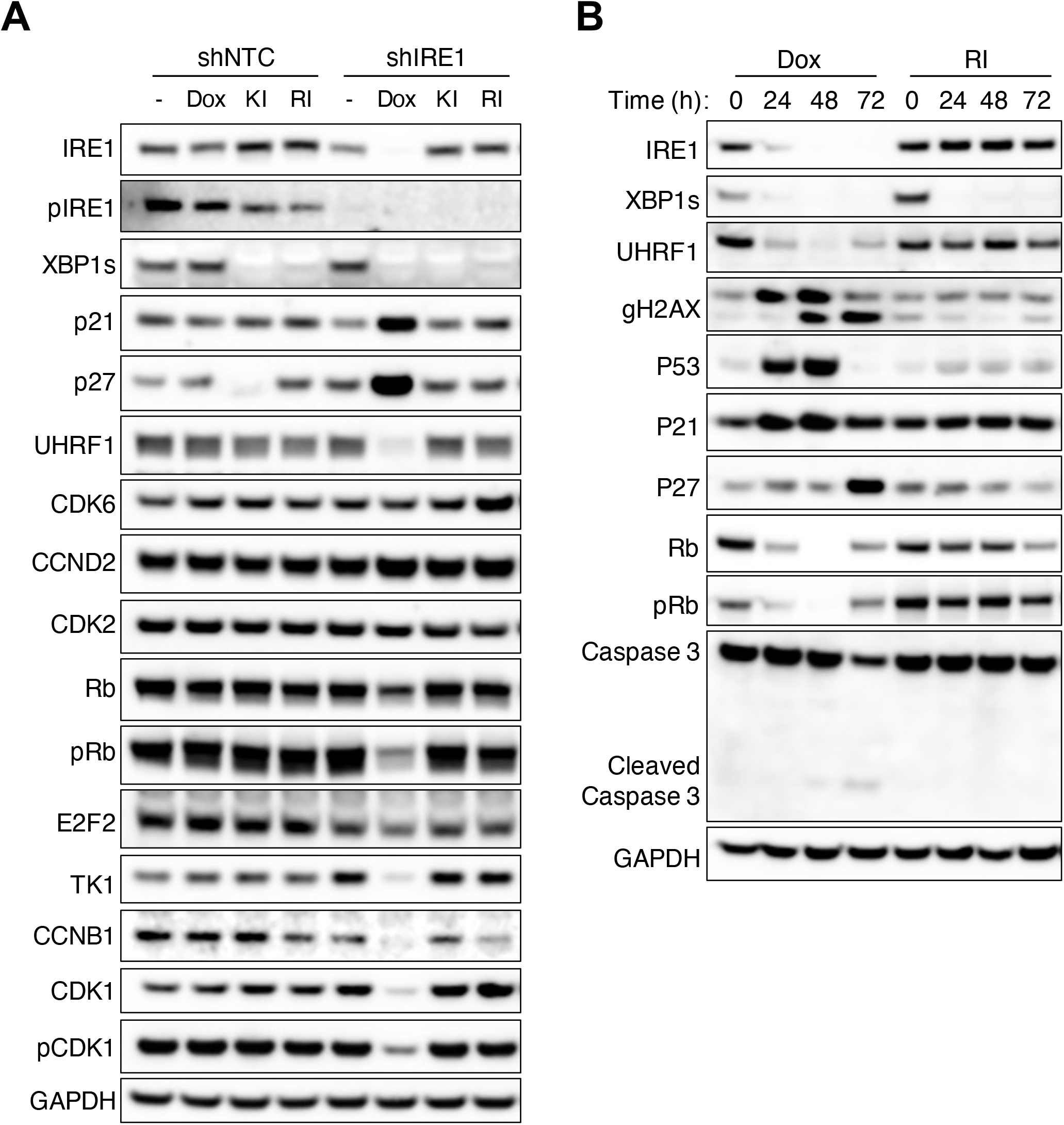
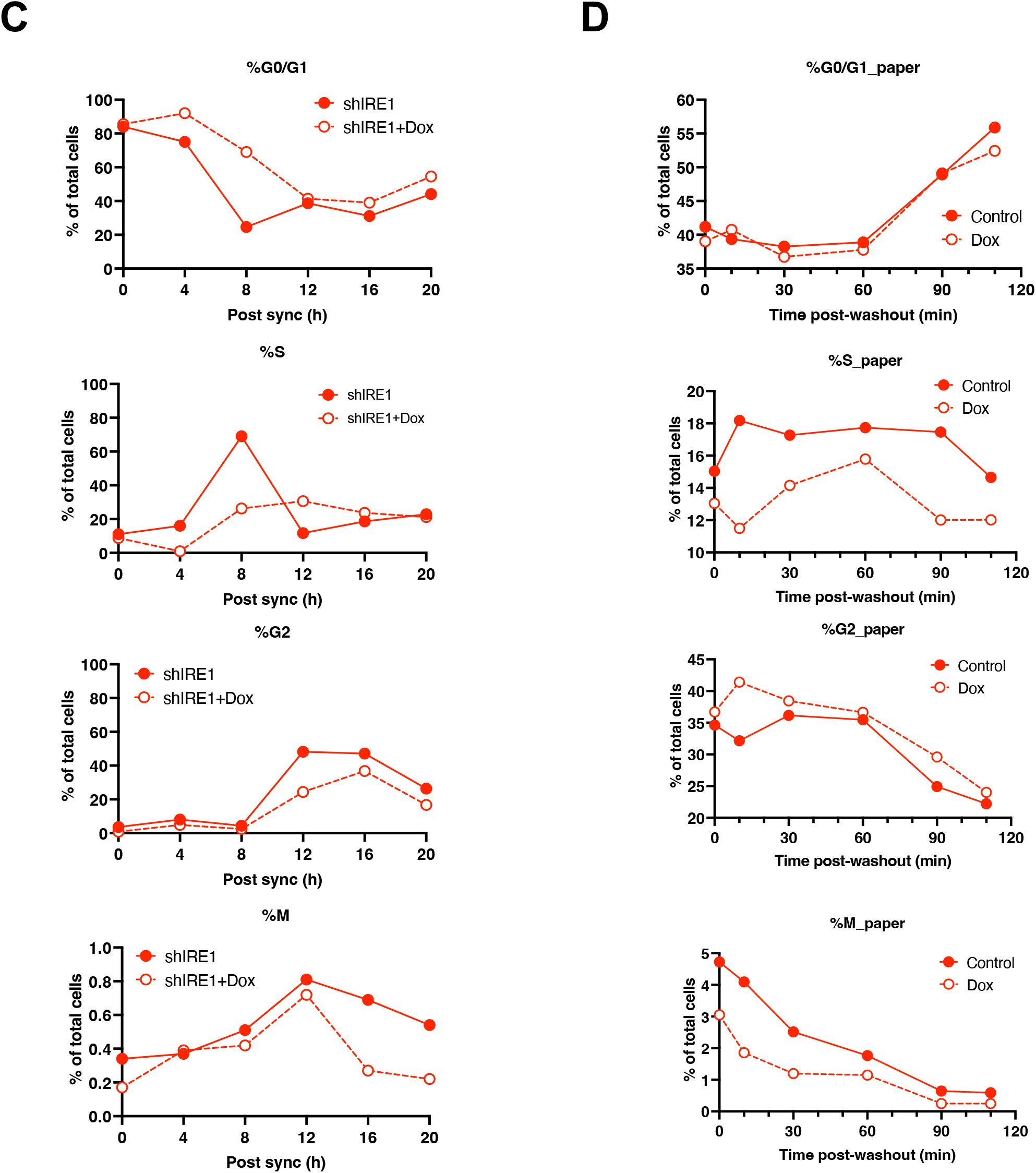
IRE1 depletion downregulates multiple cell cycle proteins. **(A)** AMO1 shNTC or shIRE1 cl.1 cells were incubated for 48 h in the absence or presence of Dox (0.2 μg /ml), or IRE1 KI or RI (3 μM) and analyzed by IB. **(B)** AMO1 shIRE1 cl.1 cells were incubated for the indicated time in the presence of Dox (0.2 μg /ml) or IRE1 RI (3 μM) and analyzed by IB. **(C)** Graphs depicting the results from Fig. 7A for each cell cycle phase. **(D)** Graphs depicting the results from Fig. 7C for each cell cycle phase.

**Figure S8.**
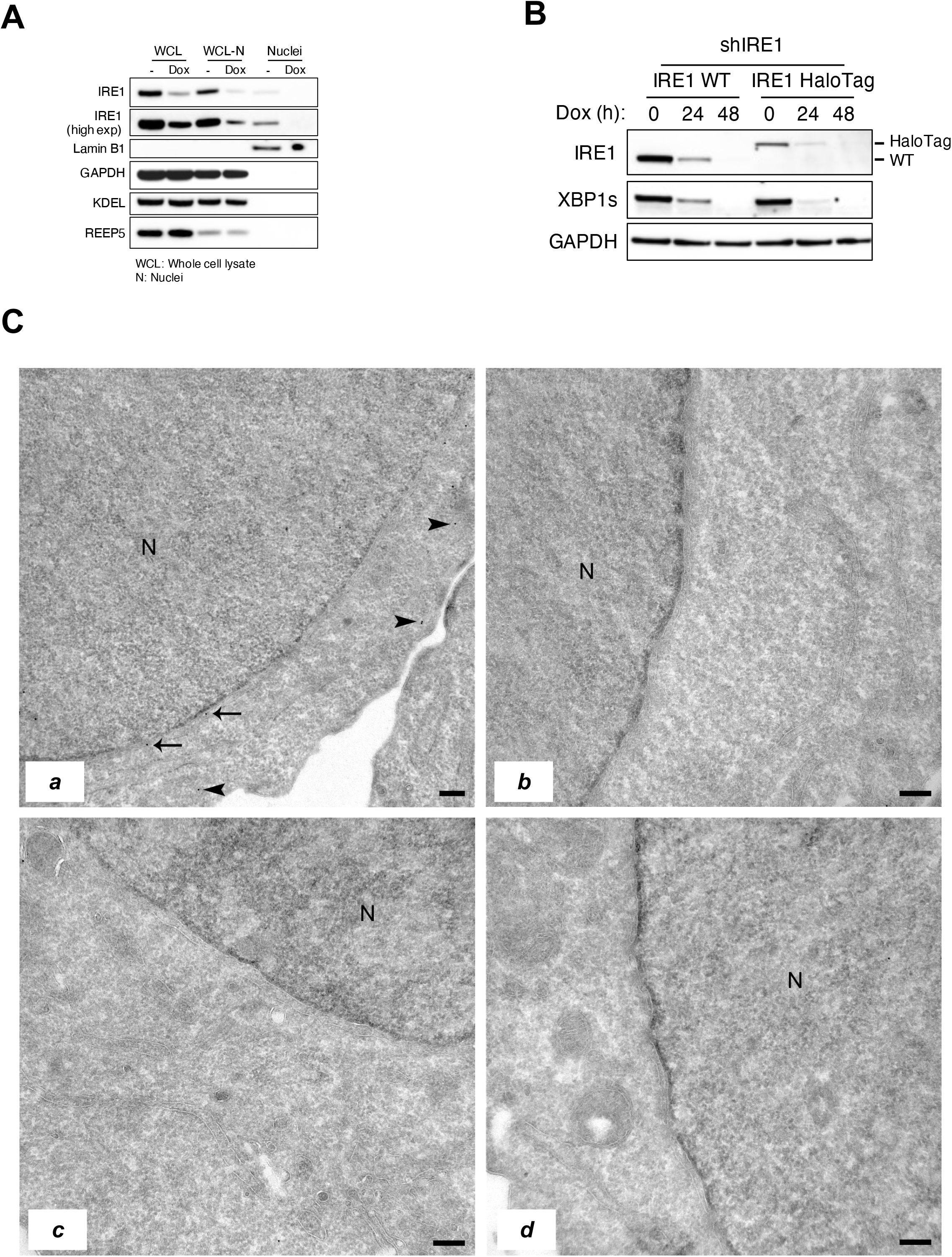
Endogenous IRE1 protein can localize to the nuclear envelope. **(A)** AMO1 shIRE1 cl.1 with wild type IRE1 (WT IRE1) or AMO1 shIRE1 cl.1 with endogenously tagged IRE1 (IRE1 HaloTag) cells used for the immuno-EM study in Fig. 8C were incubated for the indicated time in the presence of Dox (0.2 μg /ml) and analyzed by IB. Note that HaloTagged IRE1 runs slower that WT IRE1 due to the increase in IRE1 size produced by the HaloTag. **(B)** Additional examples of the staining described in Fig. 8C. **a)** Additional example of IRE1-HaloTag detection in AMO1 cells. **b)** Control staining for protein-A-Gold^10^ without anti-HaloTag antibody. **c)** Control staining for AMO1 shIRE1 cl.1 cells after 48 h of incubation with Dox (0.2 μg /ml) to deplete IRE1. **d)** Additional example as in **c)**.

## MATERIALS AND METHODS

### Cell culture and treatments

Parental AMO1, KMS27, JJN3, L363, HCT116 and OPM2 cell lines were obtained from ATCC and authenticated by short tandem repeat (STR) profiles. U-2 OS WT Flp-In T-REx and U2OS IRE1-HaloTag cells were as described elsewhere^8^. Cells were tested to ensure mycoplasma free within 3 months of use. All cell lines were cultured in RPMI1640 media supplemented with 10% (v/v) fetal bovine serum (FBS, Sigma), 2mM glutaMAX (Gibco), 100 U/ml penicillin (Gibco) and 100 μg/ml streptomycin (Gibco) in a 5% CO_2_ incubator at 37 °C.

Treatments included thapsigargin (Tg) (Sigma, 100 nM), doxycycline (Dox) (Takara, 0.2 μg/ml), pan-caspase inhibitor Q-VD-Oph (QVD) (SelleckChem, 30 μM), 5-azacitidine (5-aza) (Med Chem Express, various concentrations), CDK 4/6 inhibitor Palbociclib (Med Chem Express, 1 μM, described in Toogood et al.^65^, CDK1 inhibitor RO-3306 (Med Chem Express, 9 μM, described in Sunada et al.^66^. IRE1 kinase inhibitors KI: KI1 G5758 (described in example 176 of patent application WO/2020/056089) or KI2 G9668 (described in Guttman et al.^49^) were used at 1-3 μM and were from Genentech. IRE1 KI used in all experiments was KI1, unless otherwise specified in the figure legend. IRE1 RNase inhibitor (RI) (MKC8866) was used at 1-3 μM and was synthesized based on Sanches et al.^51^. Treatment time is indicated in the figures.

### shRNA knockdown of IRE1 or XBP1

Parental AMO1, KMS27, JJN3, L363, HCT116 and OPM2 cell lines were transfected simultaneously with three different shRNAs for either *ERN1* (IRE1) or *XBP1;* or one shRNA for non-targeting control (NTC) (see sequences below). Each shRNA was cloned in the piggyBac Dox-inducible ODE vector containing a Puromycin resistance marker. TransIT-X2 Dynamic Delivery system (Mirus) was used for transfection following manufacturer’s instructions. Transfected cells were cultured in Puromycin and surviving cells were single cell cloned. Clones with the most complete knockdown of IRE1 or XBP1 were selected by immunoblot.

- NTC shRNA: 5’ TAGATAAGCATTATAATTCCT
- IRE1 shRNA 7: 5’-AGAACAAGCTCAACTACTT-3’
- IRE1 shRNA 8: 5’-GCACGTGAATTGATAGAGA-3’
- IRE1 shRNA 9: 5’-GAGAAGATGATTGCGATGG-3’
- XBP1 shRNA 4: 5’-GGTATTGACTCTTCAGATT-3’
- XBP1 shRNA 6: 5’-GCAAGTGGTAGATTTAGAA-3’
- XBP1 shRNA 7: 5’-GCTGGAAGCCATTAATGAA-3’

### CRISPR/Cas9 knockout of Caspase-8

AMO1 shIRE1 cl.1 C8 KO cells were generated using CRISPR technology by co-transfecting a Cas9 containing plasmid, pRK-TK-Neo-Cas9, with a Caspase-8 (*CASP8*) targeting gRNA (5’-GCCTGGACTACATTCCGCAA-3’) cloned into a pLKO vector. AMO1 shIRE1 cl.1 cells were transfected using TransIT-X2 Dynamic Delivery system (Mirus), according to the manufacturer’s protocol, and single cell cloned. Clone with the most complete depletion of Caspase-8 was selected by immunoblot.

### Cell confluency, viability and caspase activity assays

Cells were plated at 5×10^3^ cells/well in ultra-low attachment (ULA) 96-well plates (7007, Corning) and centrifuged at 600g for 5 min for spheroid formation. For HCT116 cells, 2×10^3^ cells/well were seeded in a clear flat bottom 96-well plate (3595, Corning). Treatments were added at the time of cell seeding and cells were cultured in complete RPMI media to a final volume of 200 ul/well. For the conditioned media (CM) experiments, cells were instead seeded in CM collected from a confluent T-75 flask of either parental AMO1 (WT) or AMO1 shIRE1 cells treated with Dox for 24 h. The CMs were filtered with a 0.2 μm filter to remove cells and stored at -80 °C until use. For the formation of spheroids in the ULA plates, cells were centrifuged at 600 g for 5 minutes. Cultures were maintained at 37 °C throughout the duration of the experiment. All experimental conditions included at least five replicates and were repeated 3 times.

Cell confluency was tracked using a live cell imaging system (IncuCyte Zoom, Essen Bioscience). Picture frames were captured at 4-hour intervals using a 4X objective and cell confluency (%) from the total well area as a function of time was calculated using the IncuCyte software.

ATP-consumption, by Cell Titer Glo assay (Promega), and caspase activity, by Caspase-Glo 3/7 assay (Promega), were assessed using manufacturer’s instructions. Luminescence was read on an Envision system (PerkinElmer).

### Subcutaneous xenograft growth and efficacy studies

Animals were maintained in accordance with the Guide for the Care and Use of Laboratory Animals (National Research Council 2011). Genentech is an AAALAC-accredited facility and all animal activities in this research study were conducted under protocols approved by the Genentech Institutional Animal Care and Use Committee (IACUC). Mice were housed in individually ventilated cages within animal rooms maintained on a 14:10-hour, light:dark cycle. Animal rooms were temperature and humidity-controlled, between 68 to 79 °F (20.0 to 26.1 °C) and 30 to 70% respectively, with 10 to 15 room air exchanges per hour.

10×10^6^ AMO1 shIRE1 cl. 1, AMO1 shXBP1 cl. 1, KMS27 shIRE1 cl. 9 or KMS27 shXBP1 cl.13) were suspended in 1:1 HBSS and Matrigel (Corning) to a final volume of 100 μl and injected subcutaneously in the right flank of 6-8 weeks old female C.B-17 SCID mice. When tumors reached ∼150-300 mm3, mice were randomized into the following treatment groups depending on the experiment: Vehicle (5% sucrose drinking water, changed once per week); Doxycycline (0.5 mg/ml doxycycline in 5% sucrose drinking water, changed three times per week); IRE1 kinase inhibitor (KI) G5758 (250 mg/kg in 35% PEG400/55% water/10% DMSO, twice a day by oral gavage); or IRE1 RNase inhibitor (RI) (MKC8866, synthesized based on Sanches et al.^51^) (100 mg/kg in MCT formulation, twice a day by oral gavage). For tumor efficacy studies, at least 10 animals per group were treated and tumor growth was monitored until last vehicle control animal reached humane endpoint (tumor volume reached or exceeded 2000 mm^3^).

Analyses and comparisons of tumor growth were performed using a package of customized functions in R (Version 4.1.0 (2021-05-18); R Foundation for Statistical Computing; Vienna, Austria, which integrates software from open-source packages (e.g., lme4, mgcv, gamm4, multcomp, settings, and plyr) and several packages from tidyverse (e.g., magrittr, dplyr, tidyr, and ggplot2)^82^. Briefly, as tumors generally exhibit exponential growth, tumor volumes were subjected to natural log transformation before analysis. All raw tumor volume measurements less than 8 mm^3^ were judged to reflect complete tumor absence and were converted to 8 mm^3^ prior to natural log transformation. Additionally, all raw tumor volume measurements less than 16 mm^3^ were considered miniscule tumors too small to be measured accurately and were converted 16 mm^3^ prior to natural log transformation. The same generalized additive mixed model (GAMM, described in Forrest et al.^82^) was then applied to fit the temporal profile of the log-transformed tumor volumes in all study groups with regression splines and automatically generated spline bases. This approach addresses both repeated measurements from the same study subjects and moderate dropouts before the end of the study.

For pharmacodynamic studies, at least 5 animals per group were treated for 3.5 days. 6h after last oral dosing animals were sacrificed, tumors collected and protein or RNA extracted as described below.

### Immunoblot analysis

Cells were lysed or tumor tissues were mechanically disrupted using Bead Ruptor Elite (Omni) in RIPA lysis buffer (20-188, Millipore) supplemented with Halt protease and phosphatase inhibitor cocktail (ThermoFisher Scientific) and kept on ice during 30 mins. Lysates were cleared by centrifugation at 13,600 g for 15 min at 4 °C, and protein amount was analyzed by BCA protein assay (ThermoFisher Scientific). Protein was denatured by adding NuPAGE LDS buffer and DTT reducing buffer (Invitrogen) and incubating the samples at 95 °C during 5 min. Equal amounts of denatured protein were loaded in NuPAGE pre-cast gels (Invitrogen), fractioned by SDS-PAGE and electro-transferred to nitrocellulose membranes using the iBLOT2 system (Invitrogen).

Membranes were blocked in 5% nonfat milk solution for 1 hour at room temperature and probed with the corresponding primary antibody at 1:1,000 dilution overnight at 4 °C. This was followed by incubation with the corresponding horseradish peroxidase (HRP)-conjugated secondary antibody at 1:10,000 dilution during 1 hour at room temperature. All secondary antibodies were from Jackson Laboratories. The primary antibodies are listed in the Table below.

| Antigen | Reference | Species | Provider | Notes |
| --- | --- | --- | --- | --- |
| IRE1 | 3294 | rabbit | Cell Signaling |  |
| pIRE1 |  | rabbit | Genentech | Described in Harnoss et al.<sup>29</sup> |
| XBP1s |  | mouse | Genentech | Described in Harnoss et al.<sup>29</sup> |
| CD59 | ab133707 | rabbit | Abcam |  |
| Caspase-3 | 9662 | rabbit | Cell Signaling |  |
| Cleaved Caspase-3 | 9663 | rabbit | Cell Signaling |  |
| Caspase-8 | 9746 | mouse | Cell Signaling |  |
| P53 | 9282 | rabbit | Cell Signaling |  |
| pP53 | 9284 | rabbit | Cell Signaling | Phosphorylation at S15 |
| P21 | 2947 | rabbit | Cell Signaling |  |
| P27 | 2552 | rabbit | Cell Signaling |  |
| 53BP1 | 4937 | rabbit | Cell Signaling |  |
| H2AX | 7631 | rabbit | Cell Signaling |  |
| pH2AX | 05-636 | mouse | Sigma | Phosphorylation at S139 |
| UHRF1 | 21402-1-AP | rabbit | Proteintech |  |
| H3 | ab1791 | rabbit | Abcam |  |
| H3K9me3 | A-4036-050 | rabbit | Epigentek |  |
| Cyclin A2 | 4656 | mouse | Cell Signaling |  |
| Cyclin B1 | 4138 | rabbit | Cell Signaling |  |
| Cyclin D2 | 3741 | rabbit | Cell Signaling |  |
| Rb | 9309 | mouse | Cell Signaling |  |
| pRb | 8516 | rabbit | Cell Signaling |  |
| TK1 | 15691-1-AP | rabbit | Proteintech |  |
| CDC20 | 14866 | rabbit | Cell Signaling |  |
| CDCA5 | 20180 | rabbit | Cell Signaling |  |
| CDCA8 | 12465-1-AP | rabbit | Proteintech |  |
| Aurora Kinase B | 3094 | rabbit | Cell Signaling |  |
| APC3 | 12530 | rabbit | Cell Signaling |  |
| Geminin | 52508 | rabbit | Cell Signaling |  |
| Securin | 13445 | rabbit | Cell Signaling |  |
| KIF22 | ab75783 | rabbit | Abcam |  |
| CDK1 | 9116 | mouse | Cell Signaling |  |
| pCDK1 | 4539 | rabbit | Cell Signaling | Phosphorylation at Y15 |
| CDK2 | 2546 | rabbit | Cell Signaling |  |
| CDK6 | 133331 | rabbit | Cell Signaling |  |
| E2F2 | NBP2-67723 | rabbit | Novus |  |
| CDC25c | 4688 | rabbit | Cell Signaling |  |
| pCDC25c | 9527 | rabbit | Cell Signaling | Phosphorylation at T48 |
| B actin | 4970 | rabbit | Cell Signaling |  |
| GAPDH | 5174 | rabbit | Cell Signaling |  |

### RT-qPCR

RNA from tumor tissues or cells was extracted with RNeasy Plus kit (Qiagen) and quantified with a Nanodrop spectrophotometer (Thermo Fisher Scientific). Equal amounts of RNA were reverse transcribed and amplified using TaqMan™ RNA-to-C T ™ 1-Step Kit kit on the ViiA 7 Real-Time PCR System (Applied Biosystems). TaqMan gene expression assay probes (Thermo Fisher Scientific) were used to measure the expression of *ERN1* (IRE1, Hs00176385_m1), *XBP1s* (Hs03929085_g1), *DGAT2* (Hs01045913_m1), *TP53* (P53, Hs01034249_m1), *CDKN1B* (P27, Hs00153277_m1), *CDKN1A* (P21, Hs00355782_m1) and *HPRT1* (Hs02800695_m1). The delta-delta Ct (ΔΔCt) values per gene were calculated by relating each individual Ct value to its *HPRT1* housekeeping control, and then normalizing to the individual or averaged control condition. The relative quantification (RQ) was calculated as 2^-ΔΔCt^.

### Cell cycle determination by PI

1 million cells per condition were fixed in ice-cold 70% EtOH for at least 2h at 4°C. For the determination of cells in mitosis (M) inside the G2/M gate, cells were additionally stained with anti-pH3 conjugated with AF488 (06-570, Millipore) antibody diluted 1:500 in PBS-T + 3% BSA, and incubated with rocking overnight at 4°C. After washing in PBS-T + 3% BSA, samples were treated with 100 μg/ml of RNase (Zymo research) for 15 mins followed by incubation with 50 μg/ml propidium iodide (PI) for 20 mins at room temperature. Samples were run on a FACSCelesta Cell Analyzer (BD). At least 50.000 singlets per condition were analyzed for lineal PI area staining for DNA content determination. When pH3 staining was performed, pH3 area was also determined. Cell cycle distribution in G0/G1, S, G2/M and SubG1 phases was performed with FlowJo 10, using the Dean-Jett-Fox model and constraining the G2 CV to G1 CV. M population was calculated as % of cells in the G2/M gate. G2 cells alone were calculated as G2/M – M population.

### Cell proliferation assay by CFSE

AMO1 shIRE1 5-10 million cells/ml were washed twice in PBS to remove any serum and incubated with 1μM CFSE (Invitrogen) for 10 minutes at room temperature. Labeling was stopped by adding 4-5 volumes of cold complete RPMI media and incubation on ice for 3 minutes. Cells were washed thrice in complete RPMI media and 0.2ug/ml Dox was added to the treated conditions. Cells were then returned to the incubator. After 12 or 48 h, cells were harvested and 1 million cells were incubated with 1:100 LIVE/DEAD Fixable Near-IR Dead Cell Stain Kit (Life Technologies), according to manufacturer’s instructions. After washing, cells were fixed using FOXP3 Fix/Perm buffer set (BioLegend) and stored at 4 °C for a maximum of 7 days. At least 50.000 live singlets/condition were analyzed for CFSE area staining on a FACSCelesta Cell Analyzer (BD). Proliferation analysis and generation determination was performed using Modfit LT 6.0 software and the Cell Tracking Wizard Tool.

### Apoptosis detection by Annexin V and PI

Early and late apoptosis in AMO1 shIRE1 cl.1 and shXBP1 cl.1 cells previously treated with Dox was determined with the FITC Annexin V apoptosis detection kit with propidium iodide (PI) (BioLegend). Briefly, 1 million cells were stained following manufacturer’s instructions. At least 50.000 singlets were analyzed for Annexin V-AF488 and PI area staining on a FACSCelesta Cell Analyzer (BD). Percentage of cells in early (Annexin V^+^ PI^-^) or late (Annexin V^+^ PI^+^) apoptosis was determined with FlowJo 10.

### RNA sequencing

AMO1 shIRE1 cl.1, cl.3 or shXBP1 cl.1, cl.18 cells were harvested in biologic triplicates at 0, 24, 48 and 72 h after treatment with Doxycycline (Dox, 0.2 μg/ml). For bulk RNA sequencing (RNAseq), RNA from 1×10^6^ cell was first extracted with RNeasy Plus kit (Qiagen), per manufacturer’s protocol. Total RNA was quantified with Qubit RNA HS Assay Kit (Thermo Fisher Scientific) and quality was assessed using RNA ScreenTape on 4200 TapeStation (Agilent Technologies). For sequencing library generation, the Truseq Stranded mRNA kit (Illumina) was used with an input of 100 nanograms of total RNA. Libraries were quantified with Qubit dsDNA HS Assay Kit (Thermo Fisher Scientific) and the average library size was determined using D1000 ScreenTape on 4200 TapeStation (Agilent Technologies). Libraries were pooled and sequenced on NovaSeq 6000 (Illumina) to generate 30 million single-end 50-base pair reads for each sample.

### Proteomics

AMO1 shIRE1 cl.1, cl.3 or shXBP1 cl.1, cl.18 cells were harvested in biologic triplicates at 0 or 24 h after treatment with Doxycycline (Dox, 0.2 μg/ml). Samples were validated for IRE1 and XBP1s expression through immunoblotting. Protein from 1×10^6 cells was used for proteomics analysis. Cells were pelleted by centrifugation and washed with PBS then lysed in 8M urea, 50mM Tris HCl, pH 8.0 containing 1X Roche Complete Protease Inhibitor. After sonication, lysates were clarified by centrifugation and supernatants transferred. Protein quantification was performed and 50μg of each sample (plus sample pool) was digested overnight with trypsin. 1 μg of pool was analyzed (DDA) and comprised the spectral library. 1 μg of each sample was run in DIA mode (LC-MSMS) and interrogated against spectral library. Data were analyzed by both ScaffoldDIA and Spectronaut, followed by MSStats filtering.

### Micronuclei, mitotic and apoptotic cell determination by microscopy

1 million AMO1 shIRE1 cl.1 or shXBP1 cl.1 cells previously treated were harvested in 1 ml of complete media and kept at room temperature. 300ul of the cell suspension were transferred to a monolayer on a coatead cytoslide (Epredia) using the Cytospin 4 (Thermo Fisher Scientific Scientific). Sides were plunged into 4% PFA (from 16% PFA, Electron Microscopy Sciences) for fixation at room temperature during 30 minutes. Cell nuclei were counterstained with 1:2500 Hoechst 33342 (Thermo Fisher Scientific) at room temperature during 20 minutes and slides were mounted using Prolong Gold antifade mountant (Thermo Fisher Scientific) and #1.5 coverslips. Images of 10 fields/condition selected at random were acquired with a 40x objective in an Echo Revolve microscope. Micronuclei, mitotic cells and apoptotic cells were manually counted using the Cell Counter plugin in the image processing software ImageJ2 2.14.0/1.54f and normalized to the control samples.

### DNA quantification by flow cytometry

1 million AMO1 shIRE1 cl.1 or shXBP1 cl.1 previously treated cells were plated in 1ml complete media. For DNA staining, one drop of Hoechst 33342 Ready Flow (Invitrogen) was added and cells were returned to the incubator for 1h at 37 °C. Cells were then harvested, washed in PBS and incubated with 1:100 LIVE/DEAD Fixable Near-IR Dead Cell Stain Kit (Life Technologies), according to manufacturer’s instructions. At least 50.000 live singlets/condition were analyzed for Hoechst area staining on a FACSCelesta Cell Analyzer (BD). DNA quantity was determined as the Mean Fluorescence Intensity (MFI) of Hoechst with FlowJo10and normalized to the control samples.

### DNA methylation by 5-mc ELISA

Genomic DNA (gDNA) of 5 million AMO1 shIRE1 cells previously treated per condition was isolated using the AllPrep DNA/RNA Mini Kit (Qiagen). gDNA was quantified using Nanodrop spectrophotometer (Thermo Fisher Scientific) and 2-4 ul of gDNA per sample (approximately 100ng) were analyzed by MethylFlash Global DNA methylation (5-mC) ELISA Easy kit (Epigentek), following manufacturer’s protocol. Absorbance was read on an SpectraMax M2 Microplate Reader (Molecular Devices) at 450nm. Using the standard curve from the ELISA and the amount of gDNA/sample (quantified by Nanodrop Spectrophotometer from Thermo Fisher Scientific), percentage of methylated DNA (5-mC) is calculated and normalized to the control sample.

### Conventional Electron Microscopy

Conventional EM was used to visualize potential general trends in morphological changes during IRE1 or XBP1 depletion in AMO1 cells. The analysis focused on changes in nuclear morphology at 0, 24 or 48 hours after Dox addition to the cell culture. Cells were fixed in ½ Karnovsky’s fixative, postfixed in 2% osmium tetroxide, “en block” stained in 0.5% uranyl acetate and then dehydrated in an ascending series of ethanol and acetone. Embedding was in epoxy resin. After polymerization semithin sections were cut with a Leica UC7 ultramicrotome and placed on carbon coated glass slides. Sections were imaged with a Zeiss Gemini 300 scanning electron microscope and a BSD1 backscattered electron detector. Cells with high density heterochromatin (HC), euchromatin (EC), mitotic cells and apoptotic cells were manually counted. Additional details can be found in the figure legends.

### Nuclei isolation for protein expression analysis

Nuclei from 10-15 million AMO1 cells were isolated using the Nuclei EZ Prep Nuclei Isolation kit (Sigma Aldrich), according to the manufacturer’s protocol. The supernatant containing cytoplasmic components, as well as the nuclear extract were subjected to protein extraction and immunoblotting and described in the above section. Whole cell lysate (WCL) from intact cells was used as a control.

### Immunofluorescence of IRE1

1×10^5^ U2OS IRE1-HaloTag cells were grown on 24 h in Millicell EZ Slide 4-well chamber slides (Millipore), fixed in 4% PFA and permeabilized with 0.1% Triton X-100. Chambers were subjected to blocking solution (5% BSA, 0.5% gelatin, 0.05% Tween 20) during 15 minutes at room temperature, followed by overnight incubation with primary antibodies rabbit anti-HaloTag (1:500, G9281, Promega) and mouse anti-LaminB1 (0.5ug/ml, ab8982, Abcam) at 4 °C. Samples were washed and incubated with the corresponding secondary antibodies donkey anti-rabbit AF488 and donkey anti-mouse Cy3 (Jackson ImmunoResearch Laboratories) at 1:500 in blocking solution for 1h at room temperature. After washing, cell nuclei were counterstained with 1:2500 Hoechst 33342 (Thermo Fisher Scientific) at room temperature during 20 minutes, slides were detached from the chamber and mounted using Prolong Gold antifade mountant (Thermo Fisher Scientific) and #1.5 coverslips. Z-stacks of 30 sections (300nm/each) were captured with a 63x objective in a Leica SP8 confocal microscope. Leica Application Suite X (LAS X) software was used to analyze colocalization between IRE1-HaloTag and LaminB1 along the Z-stack projection. U-2 OS WT Flp-In T-REx and were used as negative control of the IRE1 staining.

### CRISPR/Cas9 endogenous tagging of IRE1 with HaloTag

Tagging of endogenous IRE1 in the c-terminus region with HaloTag in AMO1 shIRE1 cl.1 was achieved using the same methodology and reagents described elsewhere^8^. Briefly, cells were co-transfected with a plasmid encoding Cas9 with the guide RNA and a homology-directed repair (HDR) template plasmid targeted at C-terminus of *ERN1*, and single cell cloned. Clone with similar IRE1 expression than its non-tagged counterparts and with full IRE1 tagging (as demonstrated by a clear band that ran slower than WT IRE1 by immunoblot) was selected.

### Immuno-electron Microscopy

AMO1 shIRE1 cl.1 HaloTag cells were grown to 50-75% confluency in T25 flasks. They were fixed either with 4% paraformaldehyde (PFA) in 0.1 M Sorensen’s phosphate buffer (PB), pH 7.4, for 15 min at room temperature, or with 2% PFA, 0.2% glutaraldehyde (GA) in 0.1 M PB, pH 7.4, for 2 h at room temperature. The fixative solutions were replaced by 0.6% PFA in 0.1 M PB, pH 7.4, for 13 days. After rinsing in PBS and blocking in 0.15 % glycine in PBS, cells were scraped in 1% gelatin in PBS, pelleted at 800 RCF, and embedded in 12 % gelatin. Small blocks of cell pellet were cryoprotected with 2.3 M sucrose, mounted on aluminum pins and frozen in liquid nitrogen^83^. To detect IRE1-HaloTag in ultrathin cryosections of these cell pellet blocks, the G9281 rabbit anti-HaloTag antibody (Promega) was used at a dilution in the range of 1:30 to 1:500 (1:500 is the working dilution for immunofluorescence).

### Bioinformatic analysis

The bulk RNA-seq experiment was processed using the HTSeqGenie pipeline in BioConductor (Gregoire Pau, 2023). First, reads with low nucleotide qualities (70% of bases with quality<23) or matches to rRNA and adapter sequences were removed. The remaining reads were aligned to the human reference genome using GSNAP (v.2013-10-10-v2)^84^, allowing a maximum of two mismatches per 75 base sequence (parameters: ‘-M2 -n 10 -B 2 -i 1 -N 1 -w 200000 -E 1 –pairmax-rna=200000 –clip-overlap’). Transcript annotation was based on the Gencode genes database^85^. To quantify gene expression levels, the number of reads mapping unambiguously to the exons of each gene was calculated. The resulting count matrix was analysed in R (v.4.1.3; 03 November 2023) using the edgeR package (v.3.32.1)^86^. The count matrix was filtered to remove low expressed genes by keeping the features with at least 15 counts, considering library size differences, in more than the minimum number of samples of the experimental. The resulting filtered count matrix was then logtransformed and TMM-normalized with edgeR::cpm(log=T) and edgeR::calcNormFactors(method= “TMM”) to perform exploratory analyses. The matrix was further filtered to the most variable genes selected using projection score^87^ to focus on the major contributors to the variance in transcriptional state. This matrix was then scaled using base:scale() for posterior exploration. A heat map of the resulting data matrix with rows annotated by GeneOntology (GO) terms was constructed for preliminary interpretation with the function EmbolcallRNAseq::GOheatmap (https://github.com/PechuanLab/EmbolcallRNAseq;v.0.0.1). To understand the general clustering of the samples in transcriptional state, dimensionality reduction of the scaled matrix using principal component analysis was performed using PCAtools (https://github.com/kevinblighe/PCAtools; v.2.5.15). Differential expression analyses were conducted using limma (v.3.46.0)^88^. Volcano plots were produced using the package Enhanced-Volcano (https://github.com/kevinblighe/EnhancedVolcano; v.1.8.0). Gene set enrichment analysis was performed on the log-transformed fold change given by the differential expression contrasts using the GSEA function from Cluster-Profiler^89^ on the Hallmark Gene Set Collection^90^. Pathways were considered to be significant if their false-discovery-rate-adjusted P value was less than 0.2. The list of cell cycle genes used to score cell cycle activity on the transcriptomic data can be found in Kowalczyk et al., 2015^91^. PROGENy^92^ was used to score pathways of interest and obtain the TP53 pathway response genes.

### Statistics

All values are represented as arithmetic mean ± standard deviation if not otherwise indicated in the figure legends. All statistical analyses were performed using GraphPad Prism 10 (GraphPad Software, Inc.). The statistical tests are noted in each figure and significance is as follows: *, *P* < 0.05; **, *P* < 0.005; ***, *P* < 0.001; ****, *P* < 0.00001 consensus. A *p* value > 0.05 was considered non-significant (ns).

## ACKNOWLEDGEMENTS

We thank Vladislav Belyy for advice with HaloTag technology, Suzie Scales and Meredith Sagolla for coordinating EM studies. We thank Lauren Gutgesell for help with cell line characterization and Ofer Guttman, Alan Gutierrez and Bruno Alicke for assistance with in vivo models. We thank René Scriwanek for excellent assistance with photography and figure layout. The EM infrastructure of UMC Utrecht used for this work is part of the National Roadmap for Large-Scale Research Infrastructure (NEMI) financed by the Dutch Research Council (NWO), project number 184.034.014 to JK.

